# High-resolution profiling of neoantigen-specific T cell receptor activation signatures links moderate stimulation patterns to resilience and sustained tumor control

**DOI:** 10.1101/2022.09.23.508529

**Authors:** Franziska Füchsl, Johannes Untch, Vladyslav Kavaka, Sebastian Jarosch, Carolin Vogelsang, Niklas de Andrade Krätzig, Dario Gosmann, Roland Rad, Dirk Busch, Eduardo Beltrán, Eva Bräunlein, Angela M. Krackhardt

## Abstract

Neoantigen-specific T cell receptors (neoTCRs) promise a safe, highly personalized therapeutic approach in anti-tumor immunotherapy. Substantial progress has been made regarding their identification whereas detailed functional assessment of single TCR characteristics impacting therapeutic efficacy is lacking.

We previously identified and functionally characterized neoTCRs specific for neoepitopes derived from KIF2C and SYTL4 demonstrating differences in functional avidity in a patient with metastatic melanoma. In this work, we now combined single-cell TCR- and RNA-sequencing using stimulated peripheral blood derived CD8^+^ T cells of this patient and thereby identified two new neoTCRs recognizing the previously identified mutated epitope KIF2C^P13L^. Analyzing patient-derived neoTCR expressing T cells, we detected distinct activation patterns as a measure for substantial heterogeneity within oligoclonal T cell responses towards neoantigens upon specific ex vivo-restimulation. Moreover, neoTCR-transgenic T cells from healthy donors were employed for detailed in vitro and in vivo fine-characterization focusing on TCR-intrinsic functional patterns. Most importantly, in a xenogeneic mouse model experimentally simulating rechallenge of tumor infiltrating lymphocytes (TILs) after adoptive T cell transfer, we found that T cells expressing neoTCRs with a moderate activation profile provide a stable and more sustained anti-tumor response upon repeated in vivo tumor challenge as compared to neoTCRs with a stronger, burst-like reactivity. These insights have significant implications for engineering TCR-transgenic T cells for therapeutic purposes.

**One Sentence Summary:** Combining TCR specificity linked single-cell transcriptomics with in vitro and in vivo characterization of transgenic T cells helps to decipher functional potential and persistence of neoantigen-specific T cell receptors (TCRs) for TCR-transgenic T cell-based adoptive cellular anti-tumor immunotherapy.

## Introduction

Immunotherapeutic regimens have revolutionized anti-tumor therapy of multiple malignancies, especially advanced by the efficacy of immune-checkpoint inhibition (ICI) (*1*). Based on inherent immunogenicity of malignant cells, these therapies make use of the immune system’s ability to recognize and eradicate tumor cells. Though the exact interplay between immune recognition and tumor eradication versus escape during immunotherapy remains ill-defined to date, its understanding is key to develop full immunotherapeutic potential (*2*). Next to ICI, T cell-based adoptive cell therapy (ACT) approaches using tumor-infiltrating lymphocytes (TILs) or T cells genetically engineered to express T cell receptors (TCRs) or chimeric antigen receptors (CARs) have shown promising results (*3, 4*). Since one major challenge lies in attacking mutant cells with as little off-target toxicity as possible (*5, 6*), targeting neoantigens arising from somatic, tumor-restricted mutations promises a safe, precise and highly personalized approach. Targeting neoantigens with TILs or TCR-transgenic (tg) ACT confers deep, durable responses in various cancer entities (*7–9*). Prognostic biomarkers for successful immunotherapy, moreover, include tumor mutational burden, emphasizing the importance of neoantigens and neoantigen-specific T cells for anti-tumor-response (*10*). However, discovery of neoantigen-specific TCRs (neoTCRs) relied on labor-intensive functional T cell assays or sorting of reactive T cells in the past (*11, 12*). Recently, single-cell sequencing-based identification approaches led to a less biased discovery to some extent (*13–18*). Yet, low frequency in peripheral blood and dysfunction of TILs still pose major challenges for neoTCR detection (*19*).

Attempting to use neoTCRs for ACT, in-depth characterization of neoantigen-specific T cell phenotype and function as well as understanding of mechanisms and parameters affecting efficient and sustained TCR-reactivities are crucial and will shape therapeutic regimens (*20, 21*). Generally, the heterogeneous functional states of tumor-specific T cells are known to range from strong effector phenotypes to dysfunctional subsets, yet their effects onto short-and long-term tumor control remain largely unclear (*22–25*). So far, a small number of approaches combined neoTCR identification in tumor-derived TILs across different entities with transcriptomic characterization of the whole neoTCR-population although limited focus has been put on individual TCR-clonotype properties and functional patterns (*14–18*). Few preclinical models meanwhile aimed at deciphering the impact of TCR-stimulation strength onto T cells although with, however, highly limited translational significance, so far (*26, 27*). Despite attempts to transfer such results into patient datasets, translational assessment of different patient-neoTCRs for persistence of functionality in tumor settings with chronic antigen presence is missing (*27*).

In this case study, we build on previous work, where we identified mutated peptide ligands by mass spectrometry (MS) and in-silico prediction in melanoma patient Mel15. Subsequently, we investigated TILs and PBMCs from Mel15 and discovered six neoTCRs targeting the two neoantigens KIF2C^P13L^ and SYTL4^S363F^ (*12, 28*). We now combine single-cell transcriptome sequencing (scRNA-seq) and single-cell T cell receptor sequencing (scTCR-seq) and thereby identify two novel KIF2C^P13L^-specific neoTCRs. These two neoTCRs differ substantially in their transcriptomic activation profile from the previously known TCRs with KIF2C^P13L^-specificity and thereby reveal a broad functional repertoire of neoTCRs recognizing a defined common neoantigen. We comprehensively analyze common and divergent properties of neoTCR-clonotypes with specificity for a shared neoantigen. Thereby, we are not only able to show that diverse activation patterns detected in scRNA-and scTCR-seq of primary T cells are mirrored by in vitro and in vivo data of T cells transgenic for defined neoTCR indicating significant stability in structural functionality. Moreover, including a novel in vivo model for repeated tumor challenge, we also provide evidence for substantial differences in persistence of functional capacity for defined neoTCRs depending on their stimulation signatures and demonstrate that neoTCRs harboring a moderate activation pattern outperformed TCRs with burst-like reactivity upon repeated antigen encounter in vivo. These data may have an important impact on the selection and modification of tumor-reactive TCRs for their application in ACT.

## Results

### Sensitive identification of neoTCRs via scRNA-seq

We previously reported about the neoantigens SYTL4^S363F^ and KIF2C^P13L^ identified in a melanoma patient (Mel15) by our proteogenomic approach including MS as well as in silico-prediction (*12, 29*). We subsequently detected reactive T cell clones and neoTCRs derived from PBMCs or TILs of the patient with specificity for SYTL4^S363F^ or KIF2C^P13L^. Functional characterization of these neoTCRs revealed that TCR clonotypes of comparably lower avidity (KIF2C^P13L^-reactive) in comparison to those with higher functional avidity (SYTL4^S363F^-reactive) showed high frequencies within tumor, lymph node and blood of the patient and demonstrated surprisingly equal primary reactivity in vivo within a xenogeneic murine tumor model (*12, 28*).

To further understand qualitative differences in transcriptomics between these previously described neoTCRs and potentially identify novel clonotypes, we performed scTCR- and scRNA-seq on a peripheral blood sample of stage IV melanoma patient Mel15 at that time treated with Pembrolizumab in a setting of no further evidence of disease (*12*). By enriching for CD137^+^ activated T cells following specific stimulation with neoantigens SYTL4^S363F^ and KIF2C^P13L^ and employing scTCR-seq, we aimed to increase the sensitivity for detection of less frequent neoTCRs (Figure 1A). Indeed, upon antigen-specific stimulation of enriched and expanded T cells, we observed an increase in peptide-specific T cells in the enriched population with significantly up-regulated CD137 expression (Figure S1A) and increased IFN-γ secretion (Figure S1B).

**Figure 1.**
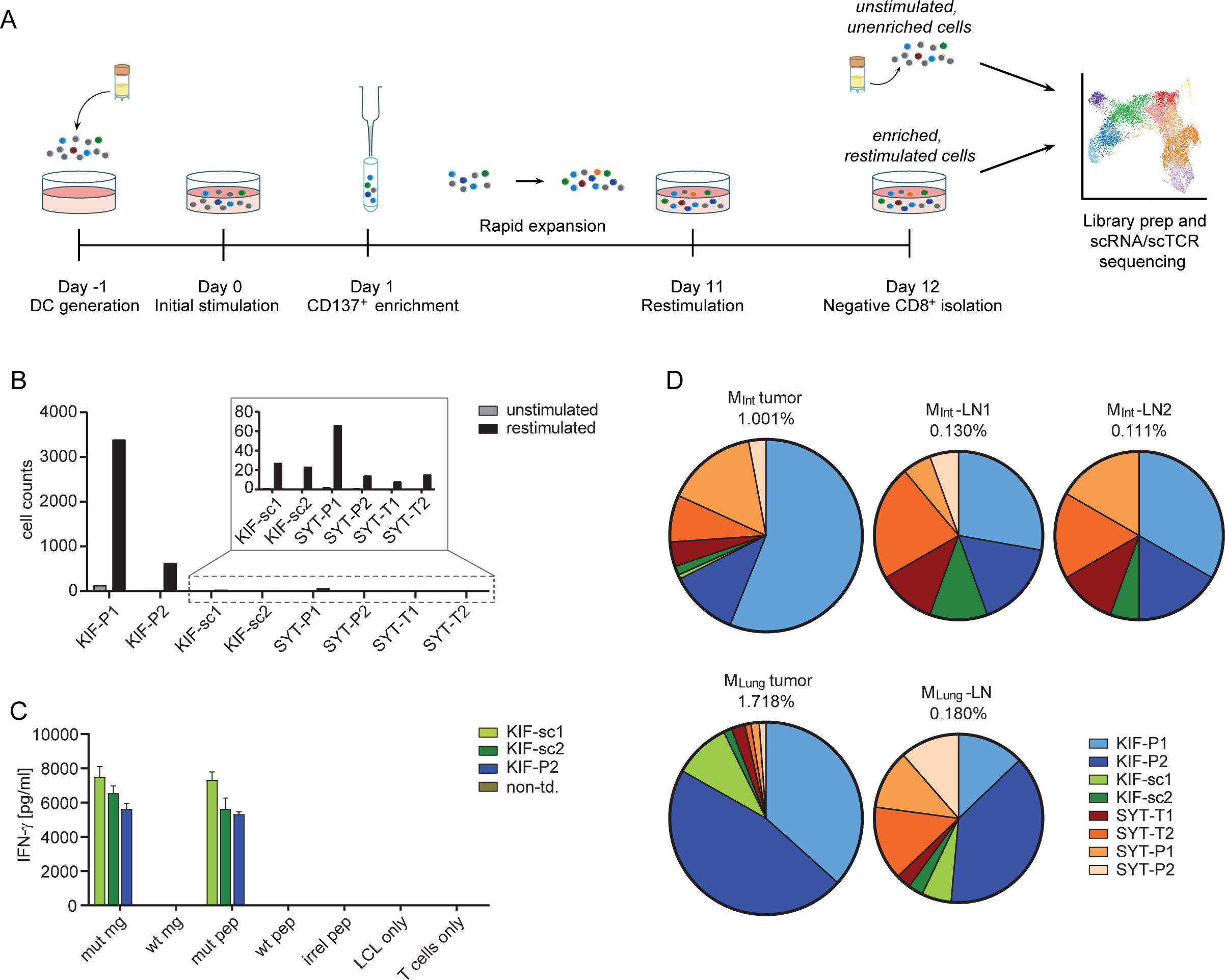
Identification of neoTCR via scTCR-seq of CD8^+^ T cells from melanoma patient Mel15. A, Schematic experiment setting. B, Increase in total number of cells of known (KIF-P1 and -P2, SYT-T1, -T2, -P1, -P2) and newly identified (KIF-sc1 and -sc2) TCRs upon antigen-specific stimulation and CD137-enrichment with dominance of KIF-P1 and -P2 harboring high precursor frequency. The two newly identified KIF2C-TCRs were selected based on fold change of TCR frequency and highest absolute frequency in the stimulated sample. C, Assessment of antigen-specific IFN-γ-secretion for the two newly identified TCRs KIF-sc1 and -sc2 in comparison to the known TCR KIF-P2. Cytokine secretion was measured by IFN-γ-ELISA upon 24h of co-culture of TCR-tg T cells from one representative donor with Mel15-LCLs transgenic for the mutated KIF2C^P13L^ minigene (mut mg) and the wildtype KIF2C minigene (wt mg) as well as pulsed for 2h at 37°C with the mutated and wildtype peptide (mut pep and wt pep). An irrelevant peptide (irr peptide), target cells (LCL only) or T cells alone (T cell only) served as negative controls. D, Frequency of KIF-sc1 and -sc2 in relation to the previously identified TCR-sequences (*28*) identified by deep sequencing of the TCR-β-chain in intestinal (MInt) and lung metastases (MLung) as well as corresponding non-malignant draining lymph nodes (MInt-LN1, MInt-LN2 and MLung-LN) of patient Mel15. Non-td: non-transduced.

The diversity of TCR clonotypes with one defined alpha and one defined beta chain in our samples decreased throughout stimulation, from 1832 different clonotypes in the unstimulated to 279 in the restimulated sample. When including clonotypes with only one defined alpha or beta chain, the numbers decreased from 2657 different clonotypes in the unstimulated to 362 clonotypes in the restimulated sample (Supplementary Table 1). All six previously known reactive receptors ranged amongst the most expanded TCRs: KIF-P1, KIF-P2, SYT-T1, SYT-T2, SYT-P1 and SYT-P2 (Figure S1C-E, Supplementary Table 2). KIF-P1 accounted for 69.0% of the restimulated clonotypes, with a high baseline of 3.2 % before enrichment and thereby greatly exceeded all other receptors in total frequency (Figure S1C, D). Overall, the enrichment of all known clonotypes suggests high efficacy of the CD137^+^-selection step included into our workflow. Besides, we sought to identify novel clonotypes with defined specificity by comparing abundance in the unstimulated and restimulated conditions (Figure 1B, Supplementary Table 1). Two of them – KIF-sc1 and KIF-sc2 – showed specific reactivity towards the mutated epitope KIF2C^P13L^ (Figure 1C). Surprisingly, their binding motives were significantly different from the previously identified KIF2C^P13L^-reactive TCRs (Figure S1F, Supplementary Table 3). In the patient, the two newly identified KIF2C^P13L^-reactive TCRs could furthermore be detected at different frequencies in several compartments: both were detectable below the previously described (*28*) high frequencies of KIF-P1 and KIF-P2 in lung and intestinal metastases as well as the respective lymph nodes (Figure 1D, Supplementary Table 4). While KIF-sc1 was less frequent in the intestinal metastasis as well as the draining lymph nodes than KIF-sc2, the opposite was true for the lung metastasis and its draining lymph node (Figure 1D, Supplementary Table 4). In conclusion, by combining CD137^+^-enrichment and frequency comparison of clonotypes identified from scTCR-seq we could enrich all previously discovered neoTCRs and identify two new neoTCRs from peripheral blood. In comparison to the previously identified, the new receptors showed significantly different binding motives as well as varying frequencies in metastases and lymph nodes.

### Negatively regulated and proliferative transcriptomic signatures in ex vivo restimulated, patient-derived T cells

Assuming a correlation between qualitative differences in T cell activation and proliferative capacity determining quantitative predominance as previously suggested (*28*), we aimed at a better understanding of the influence of cellular stimulation of different TCRs triggered by defined neoantigens. We therefore combined scTCR-seq with transcriptome analysis via scRNA-seq. Unbiased clustering using differential gene expression of the unstimulated and the CD137^+^ enriched, neoantigen-stimulated cells 24h following the second stimulus revealed eleven clusters (Figure 2A, B). As expected, unstimulated (mainly clusters 1-6, partly 7-9) and restimulated cells (mainly clusters 7-9 and partly also 5 and 6) clustered differently (Figure S2A).

**Figure 2.**
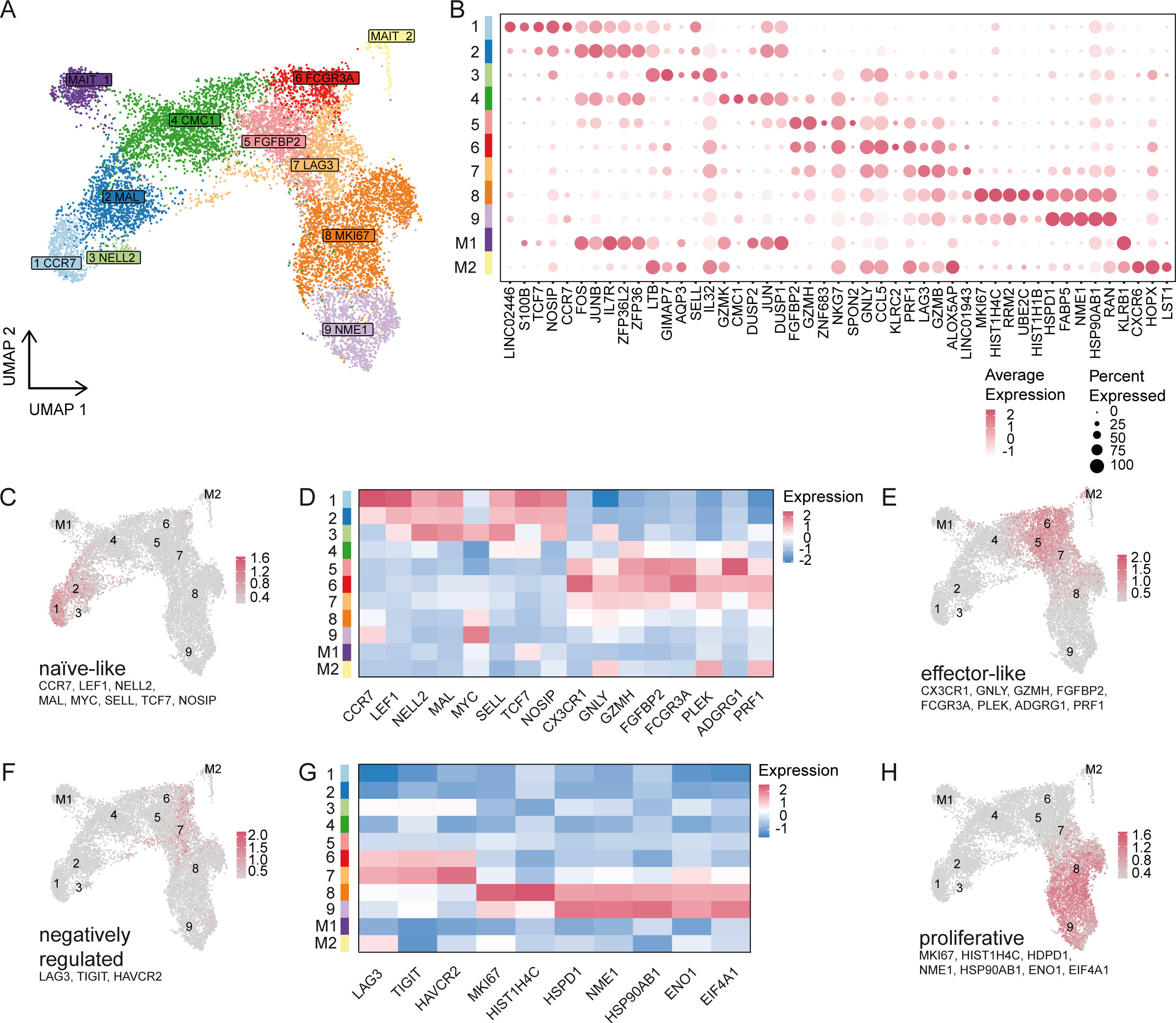
Transcriptomic profiles of ex vivo-restimulated PBMC-derived T cells stimulated with neoepitopes KIF2C^P13L^ and SYTL4^S363F^ as investigated by scRNA sequencing. A, UMAP of 5764 unstimulated and 6007 restimulated CD8^+^, sorted T cells after QC with color code indicating eleven different clusters named after one of the most differentially expressed genes each (except for MAIT1 and MAIT2). B, Dot plot showing the five most differentially expressed genes per cluster. Size of each dot indicates percentage of gene-expressing cells per cluster; color indicates scaled average fold expression of the corresponding gene within the cluster. C-E, Naïve-like/antigen-inexperienced and effector signature. UMAPs indicating distribution of naïve-like/antigen-inexperienced (C) or effector-like (E) phenotypes. Heatmap (D) showing the scaled average expression of the transcriptomic markers used for generation of signature scores as highlighted in C and E. F-H, negatively regulated and proliferative signature. UMAPs showing negatively regulated (F) and proliferative (H) cells. Heatmap of transcriptomic markers (G) integrated in signatures depicted in F and H.

We primarily focused on the distribution of T cell phenotypes across these identified clusters (Figure 2C-H). A naїve-like, antigen-inexperienced transcriptional state (expressing CCR7, LEF1, NELL) could be identified in the unstimulated sample, mainly within clusters 1, 2 and 3 (Figure 2C, D). A smaller fraction of the unstimulated as well as parts of the restimulated cells mainly clustering in 5 and 6 (partly also 8) could be assigned to an effector-like phenotype (expressing CX3CR1, GNLY, GZMH, FGFBP2, FCGR3A, PLEK, ADGRG1, PRF1), however, missing expression of proliferative genes (Figure 2D, E). Meanwhile, the upregulation of inhibitory surface receptors (most dominantly LAG3, but also TIGIT and HAVCR2) was a particular feature of clusters 6 and 7 mainly comprising stimulated cells (Figure 2F, G). Within the mainly negatively regulated, inhibitory cluster 7, pathways indicating TCR signaling and cytokine-mediated response to the cognate antigen were highly upregulated, yet proliferative processes and cell cycle G2/M-phase transition were negatively regulated (Figure S2B). In contrast, stimulated T cells in clusters 8 and 9 had vastly initiated proliferative processes upregulating typical genes such as MKI67, HIST1H4C, HSPD1, NME1, SP90AB1, ENO1, EIF4A1 (Figure 2G, H). This is prominently reflected in cell cycle phases (Figure S2C) and high numbers of features and expanded cells of these clusters (Figure S2D, E). Overall, trajectory analysis showed dynamic evolution of our cells within different states of differentiation starting at cluster 1 (most naïve) with the lowest and ending at cluster 9 with the highest pseudotime score (Figure S2F, G).

### Heterogenous gene expression patterns in neoTCRs with shared and divergent specificities

We next analyzed the distribution of all known neoTCR clonotypes within the identified clusters (Figure 2A, 3A). Regarding the cluster-related composition of different neoTCR-clonotypes, some TCRs showed marked differences in their effector, inhibitory or proliferative state (Figure 3B). From all unstimulated cells, only KIF-P1 and -P2 could be included into the comparison of cluster-distribution surpassing the subset-analysis threshold of 25 cells after quality control. Both clonotypes were mostly present in the FGFBP2-effector cluster 5 (Figure 2A, B, 3A, B). Considering the stimulated T cell population, all four KIF2C^P13L^-specific TCRs were mostly present in a proliferative state (clusters 8 and 9). Meanwhile SYT-P2, -T1 and -T2 were distinguished by a high percentage of cells from the inhibitory LAG3-cluster (cluster 7). SYT-P1 clustered more similarly to the KIF2C-pattern, raising the question about further heterogeneity within the SYTL4-TCRs. It has to be noted, however, that the absolute number of cells compared per TCR differed substantially likely associated to TCR-frequencies before stimulation among other factors (Figure 1B, Supplementary Table 2). In conclusion, these transcriptome analyses added further evidence to support the heterogeneity within activation patterns of diverse KIF2C^P13L^-and SYTL4^S363F^-specific TCRs.

**Figure 3.**
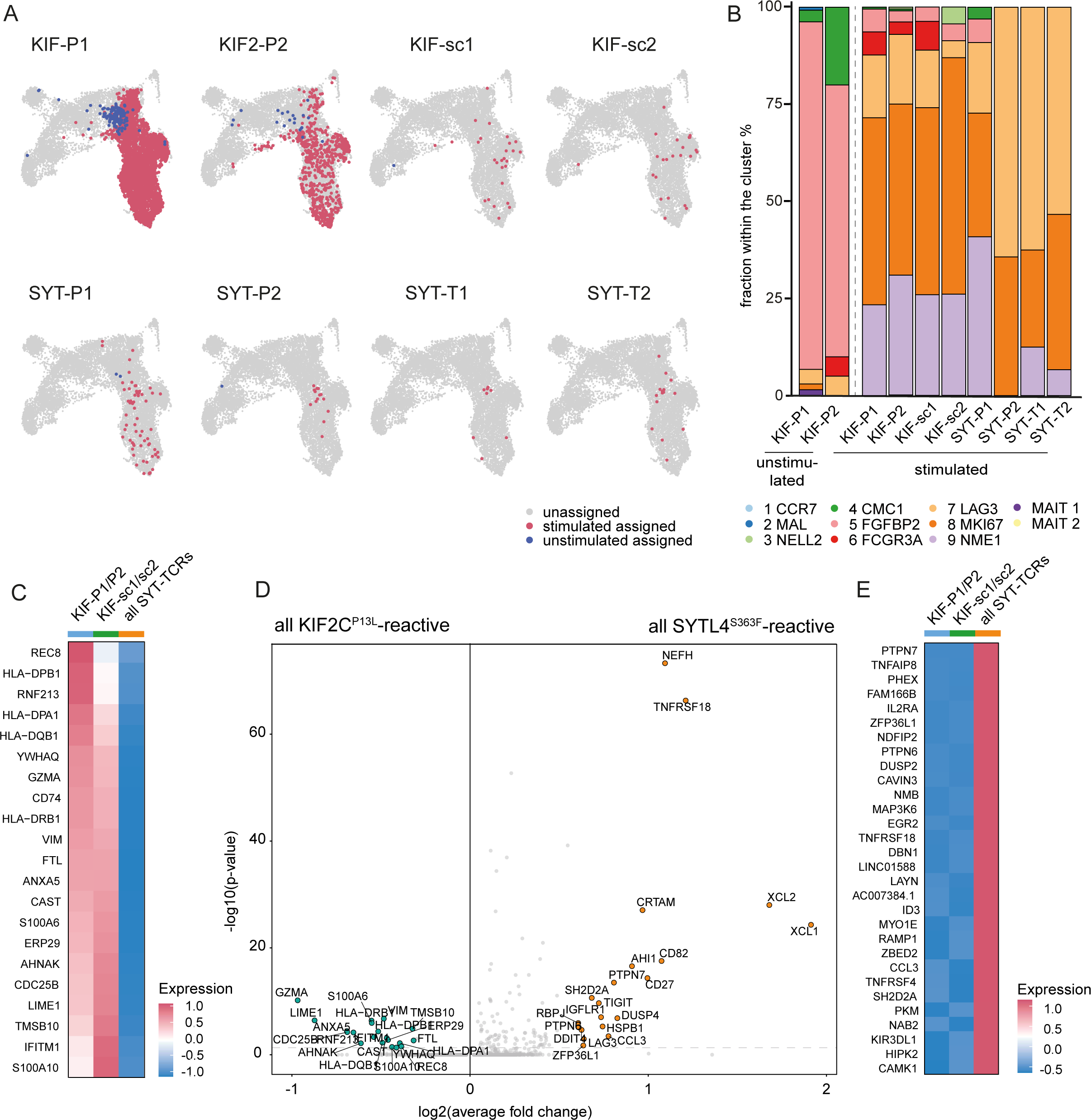
Detection of a heterogeneous spectrum of activation patterns for different neoTCRs between cytotoxic, less inhibited (KIF-P1 and -P2) and inflammation-related, but strongly inhibited activation (SYTL4^S363F^-specific TCRs). A, UMAPs of unstimulated and restimulated CD8^+^ cells showing distribution of single known TCR-specificities from both, the stimulated (red) and unstimulated (blue) sample. Non-assigned cells are depicted in grey for stimulated and unstimulated sample. B, Bar-plot indicating percentual distribution of neoTCR clonotypes per cluster. Only conditions surpassing the threshold for minimal cell numbers (>25) were included. C, Heatmap showing scaled average differential transcriptomic gene expression comparing all KIF2C^P13L^-specific neoTCRs (separating KIF-P1 and -P2 from the newly identified KIF-sc1 and -sc2) and all SYTL4^S363F^-specific neoTCRs (all SYTL4-TCRs). D, Volcano plot indicating fold changes and p values of differential transcriptomic gene expression comparing all KIF2C^P13L^- (all previously and all newly identified TCRs) and SYTL4^S363F^-specific TCRs. E, Heatmap showing scaled average differential transcriptomic gene expression comparing the same neoTCR-groups as in C, ranked with focus on highest expressed genes in SYTL4^S363F^- specific TCRs.

An unbiased look at the differentially expressed genes of KIF2C^P13L^- versus SYTL4^S363F^ -specific TCRs reflected the patterns described. On the one hand, KIF2C^P13L^-specific TCRs upregulated genes of cytotoxic effector functions (GZMA), antigen presentation (MHC class II-genes and CD74) and TCR-signaling genes (ANXA5, AHNAK, S100A6, S100A10, LIME1) among which many are involved in calcium dependent processes (Figure 3C, D). SYTL4^S363F^-specific TCRs, on the other hand, highly expressed genes correlated with chemokine profiles and proinflammatory pathways (e.g., XCL1, XCL2, CD27, CCR3, CCL3). Thus, SYTL4^S363F^- and KIF2C^P13L^-specific TCRs showed divergent effector pathway regulation upon stimulation (Figure 3D, E, S3A, B). At the same time, inhibitory receptors like LAG3 and TIGIT (potentially also TNFRSF18), but also DUSP4 and PTPN7, two MAP-Kinase inhibitors, were upregulated in SYTL4^S363F^-TCRs implicating simultaneous inhibitory counterregulation. In contrast, the significantly upregulated genes for KIF2C^P13L^-TCRs did comparatively not include markers of inhibitory signaling (Figure 3C, D). Looking deeper into the diverse KIF2C^P13L^-specific TCRs, differential gene expression reveals a distinct state for KIF-sc1 and -sc2 differing from the described qualitatively contrasting activation signatures. This is displayed by a gradient detectable in the expression level of MHC class II genes, CD74 and GZMA from KIF-P1 and -P2 over -sc1 and -sc2 towards SYTL4^S363F^ - TCRs (Figures 3C). The heterogeneity between KIF2C^P13L^-specific profiles was further supported by the direct comparison of KIF-sc1/-sc2 versus SYTL4^S363F^-TCRs showing only the upregulation of genes associated with TCR signaling like LIME1 and S100A10 in KIF-sc1/-sc2 in contrast to upregulation of negative regulators like PTPN7 and DUSP4 only in SYTL4^S363F^-TCRs (Figure S3C). MHC class II genes, however, were not differentially upregulated between KIF-sc1/-sc2 and SYTL4^S363F^-TCRs (Figure S3C). Regarding unbiased analysis of unstimulated neoTCRs, again only KIF-P1 and -P2 transcriptomes could be analyzed comprising sufficient cell counts. Comparing both TCR clonotypes with all other unstimulated T cell clones, cytotoxic markers including FGFBP2, GZMB, GZMH, GNLY and NKG7 were predominantly upregulated (Figure S3D).

Overall, we describe a spectrum of TCR-dependent T cell activation patterns in this scRNA-seq dataset from an ex vivo restimulation setting from patient-derived neoantigen-specific CD8^+^ T cells. We detected cytotoxic, proliferative and especially less inhibited T cell activation patterns for KIF-P1 and -P2 and comparatively higher expression of inflammation- and chemokine-related as well as inhibitory genes for SYTL4^S363F^-specific TCRs. KIF-sc1 and -sc2 showed features of both patterns.

### Inflammation-related, inhibitory neoTCR-transcriptome signatures correspond to functional patterns of activation and negative counterregulation

All cells originating from patient PBMCs possess a certain differentiation state at the time of blood collection due to multiple variables including potential previous encounter with their cognate antigen as well as therapeutic regimens. To circumvent potential bias between T cell populations with different previous fates in the patient, we further compared the different neoTCRs after retroviral transduction into activated CD8^+^ T cells in independent experiments using several healthy donors. This in vitro analysis enabled antigen dose-titrated T cell stimulation and moreover helped decouple TCR-intrinsic features from patient-specific cellular differentiation.

Next to the two newly identified TCRs KIF-sc1 and -sc2, we selected TCRs SYT-T1 and KIF-P2 for these analyses covering the whole spectrum of determined functional avidities (*28*).

We investigated the effect of different stimulation strengths by analyzing functional reactivity against target cells pulsed with varying peptide concentrations (Figure 4A-C). This clearly illustrated stronger, more sensitive activation of SYT-T1-tg T cells after 24h reflected by IFN-γ secretion compared to KIF-P2 as previously described (*28*). KIF-sc1 and -sc2 showed intermediate effector functions between those two diverse reactivity patterns with KIF-sc1 ranging closer to SYT-T1, and KIF-sc2 to KIF-P2 (Figure 4A). EC_50_-value measurement for all four KIF2C^P13L^-specific TCRs from five donors confirmed slightly higher functional avidity for the two newly identified TCRs compared to KIF-P1 and -P2 independent from transduction rates and selected donor (Figure S4A, Supplementary Table 5). These differences in activation capacity were similarly reflected by surface staining of the activation marker CD137 (Figure 4B) as well as the inhibitory receptor PD-1 (Figure 4C).

**Figure 4.**
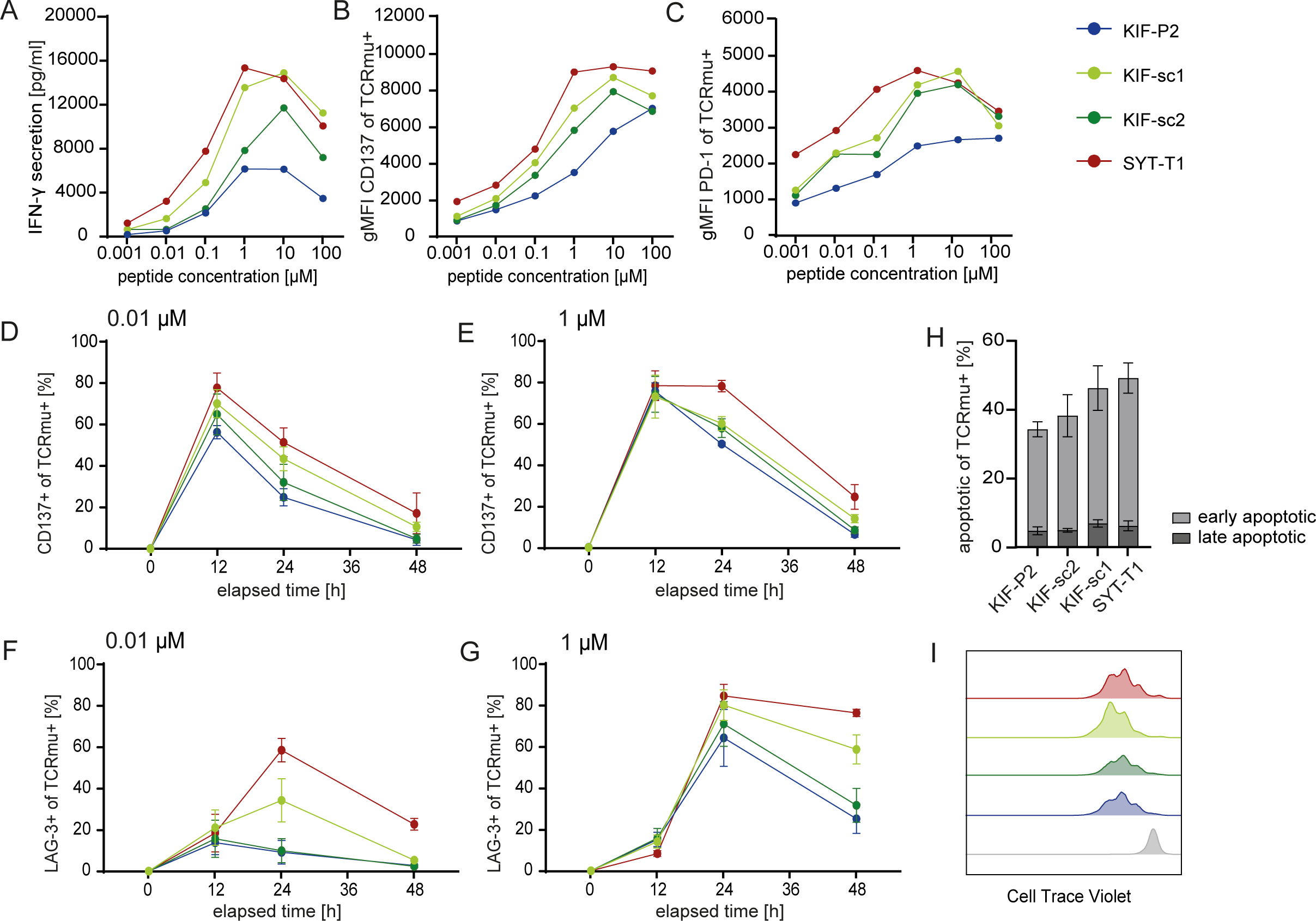
Activation strength of CD8 neoTCR-tg T cells from healthy donors is associated with inhibitory counterregulation and early apoptosis during co-cultures with different cell lines. A-C, Mel15 LCLs were pulsed with titrated peptide concentrations (2h, 37°C) and co-incubated with TCR-tg T cells with subsequent ELISA-based assessment of IFN-γ-secretion within 24h of co-culture (A). The cellular activation level was determined after 24h by FACS staining of the extracellular level of CD137 (B) and PD-1 (C) expression (reflected by geometric mean of all CD3^+^CD8^+^/TCRmu^+^ cells). The mean for ELISA data is depicted for technical triplicates of one donor; triplicates from the same donor have been pooled prior to EC FACS-staining. E:T = 1:1 (15.000 tg T cells:15.000 tumor cells). D-G, EC FACS staining at different timepoints after co-culture setup displays temporal dynamics of T cell activation marker CD137 (D, E) and inhibitory receptor LAG-3 (F, G) for TCR-tg T cells upon co-culture with JJN3-B27 peptide-pulsed target cells. A weak (0.01 µM for peptide pulsing; D, F) versus a strong (1 µM for peptide pulsing; E, G) stimulus were compared. E:T = 1:1 (10.000 tg T cells:10.000 tumor cells). H, Annexin-V/PI-staining was employed for detection of activation induced cell death (AICD) after 20h of co-culture upon strong stimulation with 1µM mut-peptide pulsed Mel15 LCLs (early apoptotic = AnnexinV^+^PI^-^, late apoptotic = AnnexinV^+^PI^+^). E:T = 1:1 (30.000 tg T cells:30.000 tumor cells). I, Representative FACS plot of a healthy donor of CTV-analysis for all TCRmu^+^ cells depicted after 4 days of co-culture with 1µM mut-peptide pulsed Mel15 LCLs (colors were chosen according to Figure B-E; representative wt mg-control depicted in grey). E:T = 1:1 (30.000 tg T cells:30.000 tumor cells). For all co-cultures in D-I technical triplicates per donor were pooled prior to staining; the mean and SD for biological replicates from three different human donors are shown.

We next analyzed temporal dynamics to decipher potential differences in kinetics of activation maxima. Performing surface staining of CD137 over the course of 48h on different TCR-tg populations upon co-culture with two different strengths of stimuli (0.01 μM and 1 μM peptide-pulsing), maximal expression of the activation marker could be detected consistently after 12h irrespective of TCR, healthy donor or stimulus with SYT-T1 again demonstrating the maximal and KIF-P2 the minimal functional reactivity (Figure 4D, E). The strong compared to the weak stimulus, nevertheless, prolonged the time of CD137 expression on a population level for all TCRs and moreover increased CD137^+^ fractions particularly for the KIF2C^P13L^-reactive TCRs. The same pattern was detected when stimulating TCR-tg T cells with another tumor cell line (Figure S4B, C).

Based on the inhibitory signature detected for SYTL4^S363F^-specific TCRs in the transcriptome analysis, we also investigated inhibitory receptors typically upregulated throughout early T cell activation. Choosing LAG3 as the most prevalent gene in the inhibitory cluster 7 (Figure 2A, F), we detected maximal upregulation of this marker after 24h especially on SYT-T1- and KIF-sc1-tg T cells as well as clear dependence on the strength of stimulus (Figure 4F, G). Most interestingly, KIF-P2 and KIF-sc2 initiated only very low LAG-3 upregulation upon the weak stimulus with a very low maximum already at 12h followed by immediate downregulation. However, upon the strong stimulus higher expression of LAG-3 similar to the other receptors was detected with a maximum at 24h for both TCRs (Figure 4F, G). Similar trends could also be shown for PD-1 levels despite overall lower expression compared to LAG-3 (Figure S4D, E). In addition, the stronger TCR activation pattern of SYT-T1 and KIF-sc1 was linked to an increased percentage of apoptotic (Annexin V^+^) cells in co-culture with diverse cell lines pulsed with 1 μM peptide (Figure 4H, S4F- G).

We continued comparing proliferation of transgenic T cells within the first few days after stimulation. On day 4 of co-culture, only slight differences between all TCR-tg T cells with a trend towards stronger proliferation of SYT-T1 and KIF-sc1 were detectable (Figure 4I, S4H-J). Thus, despite highly expressed counterregulatory receptors, proliferative dysregulation did not appear to be a key feature of TCR-tg T cells with strong activation patterns upon this first in vitro stimulation.

In summary, KIF-sc1-tg T cells show patterns more similar to SYTL4^S363F^-specific TCRs, while KIF-sc2 showed less pronounced counterregulation and remained closer to the KIF-P2-pattern. This stressed a broader heterogeneity of activation patterns even within one peptide-HLA-specificity. Overall, T cells transgenically expressing TCRs associated with proinflammatory, yet negatively regulated transcriptomic signatures, performed more sensitive. They reached higher overall levels of cytokine secretion, activation markers, but also increased inhibitory receptor expression upon first antigen encounter in vitro. This highly responsive, burst-like pattern could be described for SYT-T1 and partly KIF-sc1. In comparison, KIF-P2 and KIF-sc2 appeared with a more moderate activation signature.

### Highly responsive, burst-like TCR-signal associates with deteriorated tumor control upon repeated neoantigen challenge in vivo

To assess functionality in vivo, we investigated the potency of neoTCR-tg T cell populations including the novel clonotypes in a previously established in vivo xenograft model with the HLA-matched B cell lymphoma cell line U698M expressing minigenes coding for KIF2C^P13L^ and SYTL4^S363F^ (mut mg). The initial model designed for highest efficacy in tumor rejection (Figure 5A) revealed comparably potent rejection kinetics for all neoTCRs compared to an irrelevant, MPO-specific TCR-control (2.5D6) (*28*). We have already published the data from this experiment for KIF-P2, SYT-T1 and 2.5D6 (*28*) and now additionally show the data for KIF-sc1 and -sc2 measured throughout the same experiment. In this setting, the two newly identified neoTCRs performed equally well compared to those previously known and reached complete tumor rejection in all mice with significantly prolonged survival (Figure 5B, C).

**Figure 5.**
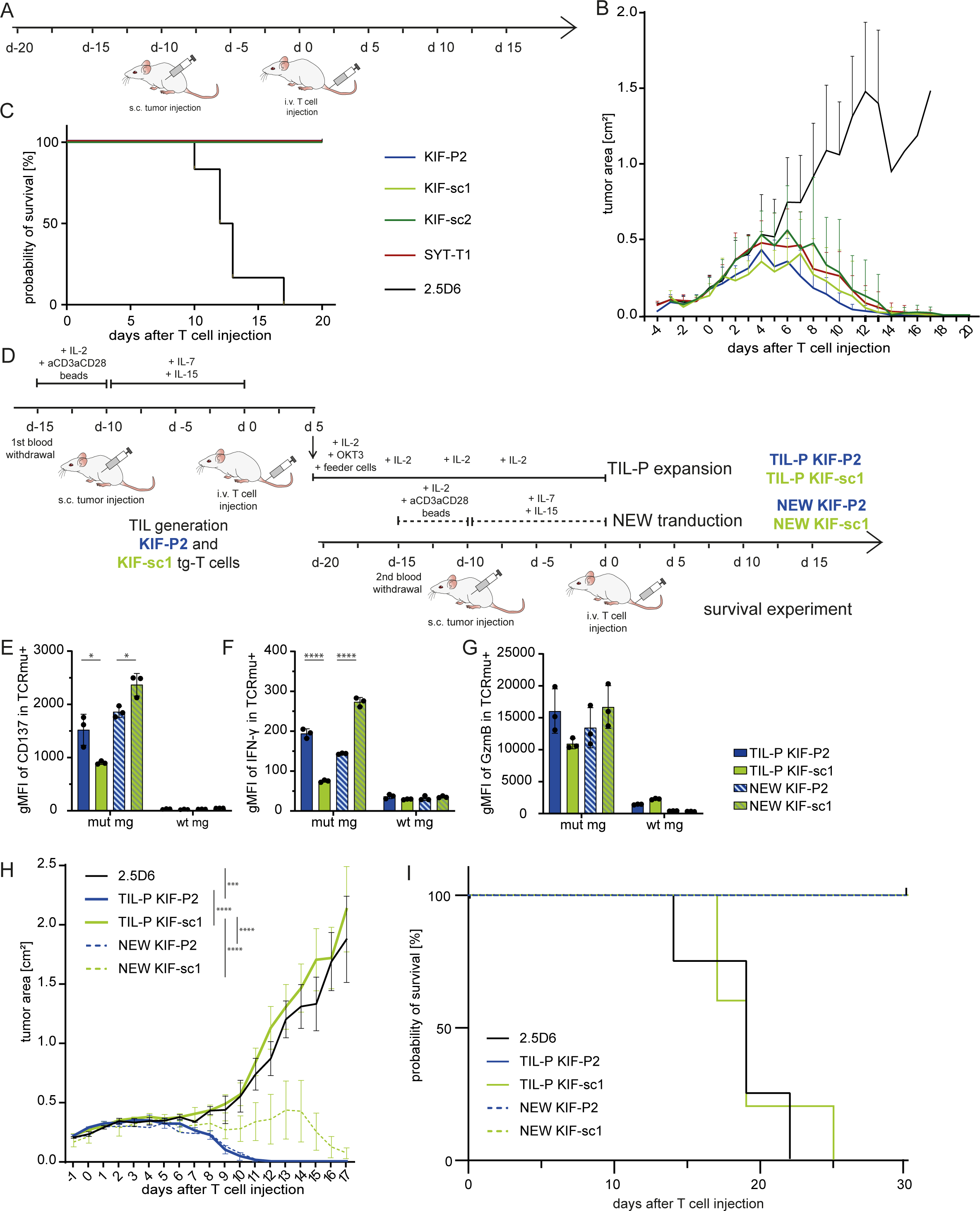
Moderate activation pattern of KIF-P2 T cells associates to sustained anti-tumor response upon in vivo rechallenge in contrast to burst-like activated KIF-sc1. A, Schematic experiment setting of xenograft neoTCR-tumor rejection experiment with newly transduced T cells. B, Tumor growth kinetics are displayed as tumor area (in cm^2^) for mut mg-U698M-tumor-bearing NSG-mice comparing neoTCR-tg T cells to the irrelevant TCR 2.5D6 until day 20. Mean values and SDs for each group of mice display rejection dynamics (n=6). Parts of this dataset were already published before (*28*). In this renewed version, tumor rejection kinetics of KIF-sc1 and -sc2 analyzed in the same experiment are included. C, Kaplan-Meier-survival curve is displayed until day 20 for tumor-bearing mice injected with different neoTCR-tg T cells. D, schematic experiment setting of xenograft neoTCR-TIL-rechallenge experiment. E-G, Ex vivo restimulation of T cells derived from TIL products on day 21 after tumor explant (TIL-P) compared to newly transduced (NEW) TCR-tg T cells from the same human donor stained for CD137 (EC; E), IFN-γ (IC; F) and GzmB (IC; G); expression was analyzed using geometric mean of all CD3^+^CD8^+^/TCRmu^+^ cells after 18h of co-culture. Mut-mg and wt-mg U698M cells were used as target cells in E:T = 1:1 (50.000 tg T cells:50.000 tumor cells). Mean and SD are shown for three experimental replicates. Statistical significance is calculated with one-way ANOVA and Tukey’s multiple comparison test (*p≤0.05, ****p≤0.0001). H, Tumor growth kinetics are displayed as tumor area (in cm^2^) for NSG-mice continuing the experiment until day 17 after second injection of in total 5x10^6^ neoTCR-tg T cells (transduction rate of 55% equalized for all groups). For the TIL- P-groups, TIL-P from two mice per TCR (highest numbers of expanded neoTCR-tg T cells) were pooled. Mean values and SEMs for each group of mice display rejection dynamics (n=5 for experimental groups, n=3 for 2.5D6; due to achievement of humane endpoint criteria one 2.5D6- control mouse was sacrificed on day 13 and excluded from this graph). Statistical significance is calculated for the tumor area on day 17 with one-way ANOVA and Tukey’s multiple comparison test (***p≤0.001, ****p≤0.0001). **I**, Kaplan-Meier-survival curve is displayed for tumor-bearing mice injected with different TCR-tg T cells (n=5 for experimental groups, n=4 for 2.5D6 control). Survival of mice receiving TIL-P-KIF-P2 compared to TIL-P-KIF-sc1 was significantly prolonged (p=0.0019, Mantel-Cox test).

Subsequently, we focused closely on two TCRs associated to either the moderate (KIF-P2) or the burst-like (KIF-sc1) activation patterns based on the novel in vitro data while sharing neoantigen-specificity and HLA-restriction. In a new experiment setting (Figure 5D) we continued challenging the in vivo model by lowering its effector cell saturation to a quarter of TCR-tg T cells. Upon sacrifice on day 5 after adoptive T cell-transfer, no measurable differences in tumor growth were observed as expected (Figure S5A). Ex vivo, we detected trends of higher IFN-γ and CD137 expression within KIF-sc1- as compared to KIF-P2-tg T cells (Figure S5B, C).

Following up the inhibitory patterns upon antigen encounter in vitro on burst-like TCR-tg T cells, we aimed to investigate repeated in vivo tumor challenge. Therefore, we generated TIL products (TIL-P) from these mice and after ex vivo expansion reinjected these cells into new tumor-bearing recipients (TIL-P were pooled from two mice per TCR). In parallel, we performed a new transduction with the same two TCRs on CD8^+^ T cells from the same donor (NEW) as control groups (Figure 5D).

Prior to reinfusion of TIL-P, we performed ex vivo co-cultures with the mut mg-tg U698M tumor cell line to compare T cell activation patterns. As expected, the newly transduced cells (NEW) proved higher activation among KIF-sc1-tg T cells upon antigen-specific stimulation in vitro as compared to KIF-P2-tg T cells (Figure 5E-G). In contrast, we detected lower levels of activation markers and effector cytokines for TIL-P-KIF-sc1-tg T cells compared to TIL-P-KIF-P2-tg T cells suggesting lower reactivity upon antigen rechallenge (Figure 5E-G). Moreover, TIL-P-KIF-sc1-tg cells showed deteriorated proliferation compared to TIL-P-KIF-P2-tg T cells ex vivo (Figure S5D). This difference was not detected for the two NEW TCR-tg T cell populations (Figure S5E).

Subsequently, we injected TIL-P containing TIL-P-KIF-sc1-tg or TIL-P-KIF-P2-tg as well as NEW TCR-tg T cells into U698M (mut mg)-tumor-bearing mice and measured tumor growth for 30 days in comparison to NEW-control groups (Figure 5D). While the TCRs of the newly transduced TCR-tg T cells of both TCRs reached complete tumor rejection in all mice until day 18 as previously described, we observed clear dysfunction of TIL-P-KIF-sc1 upon rechallenge in vivo (Figure 5H). Tumors in TIL-P-KIF-sc1 mice could not be controlled by the applied T cell-product as observed for animals receiving the non-specific T cell product (TCR 2.5D6). TIL-P-KIF-P2, meanwhile, reached complete tumor rejection in all mice (at least up to day 30) and performed equally efficient compared to the newly transduced T cells (NEW) (Figure 5H, I; S5F-H). Based on these data we conclude functional impairment for higher avidity, more burst-like activated TIL-P-KIF-sc1-tg T cells in antigen-specific T cell rechallenge and subsequently anti-tumor activity upon restimulation in vivo.

## Discussion

To date, efficacy of ICI and defined adoptive T cell therapies using TILs are assumed to rely substantially on CD8^+^ cytotoxic T cell responses, recognizing neoantigens presented on the HLA- complexes of tumor cells (*22, 30–34*). Neoantigen-reactive TCR-clones typically represent minor fractions among TILs and comprise scarce populations in human blood (*14, 19, 28*). Identification and characterization of neoTCRs, therefore, still pose a major bottleneck for selecting T cells and TCRs with favorable characteristics for effective ACT. Several approaches already aimed at enrichment of tumor-reactive T cells, exemplarily by sorting for CD137^+^ or PD-1^+^ T cells (*11, 35*). In contrast to other recent studies on TIL-derived neoTCRs (*14, 16, 36*), we used peripheral blood-derived T cells of a metastatic melanoma patient under ICI treatment with known neoantigen-specific T cell reactivity and present a restimulation-dependent single-cell sequencing approach for identification of neoTCRs and subsequent in-depth fine-characterization of these TCRs in vitro and in vivo.

The sequential approach of specific stimulation with MS-approved epitopes (*12, 29*), magnetic enrichment of CD137^+^ cells and in vitro restimulation enabled the sensitive detection of T cell clones specific for the two known neoantigens SYTL4^S363F^ and KIF2C^P13L^ despite partially very low precursor frequencies. Beyond detection of all six previously described neoTCRs (*12, 28*), two additional neoantigen-reactive TCRs with specificity for KIF2C^P13L^ were identified. This suggests peripheral blood as a valuable, easily accessible source for detection of strong neoTCRs possibly outperforming the identification of exhausted and dysfunctional neoantigen-specific TILs from tumor material. (*14, 37, 38*). In fact, dysfunction markers such as CXCL13, CD39 or CD69 have been proposed as bio- or selection markers for neoantigen-specific TILs with potential for diagnostic or therapeutic exploitation (*14–18*). However, we did not observe notable transcriptomic upregulation of such markers among patient-derived, non-restimulated KIF2C^P13L^-specific T cells potentially due to analysis of peripheral blood instead of TILs.

Since T cell effector functions are defined by distinct activation properties, we went beyond a static signature of patient-derived tumor-specific T cells (*14–18*) by specifically restimulating peripheral blood-derived T cells of Mel15 to decipher TCR-intrinsic activation patterns. Of note, upon restimulation we observed a heterogeneous pattern in neoTCR-dependent transcriptomics of these patient-derived T cells revealing qualitative differences between the identified neoTCRs. On one end of the observed spectrum, we identified features of the in synopsis with analyses on TCR-tg cells defined burst-like, highly responsive but simultaneously strongly inhibited activation patterns associated with SYTL4^S363F^-specific T cells harboring slightly higher functional avidity (*28*). These cells were characterized by strong upregulation of proinflammatory markers and chemokines, e. g., the inflammatory chemokines XCL1 and XCL2, both regularly expressed by natural killer cells and activated CD8^+^ T cells (*39, 40*). Simultaneously, these cells also significantly upregulated inhibitory receptors (LAG3, TIGIT, HAVCR2) throughout the first 24h of stimulation. Furthermore, SYTL4^S363F^-specific T cells showed upregulation of DUSP4 and PTPN7, both negative regulators of the mitogen activated protein kinase (MAPK) (*41–43*). Interestingly, these neoantigen-specific T cells were found at comparably low frequencies in the patient pointing towards an association of a very high TCR-signal strength with defects in MAPK phosphorylation and subsequent proliferative defects as previously described (*44*). On the other end, KIF-P1 and -P2, neoTCRs with overall slightly lower functional avidity, but comparably high frequencies in the patient, demonstrated a distinct, in conjunction with functional data later defined moderate activation pattern with lower opposing negative regulation. The marked transcriptomic upregulation of GZMA suggested cytotoxic capacity (*45*), while the presentation of HLA-class II molecules and CD74 may be associated to T cell-mediated antigen-presentation and proliferation (*46–48*). The expression of genes related to calcium-dependent TCR-signaling such as ANXA5 (*49*), AHNAK (*50*), S100A6, S100A10 (S100 calcium binding proteins) (*51*) and with a lesser extent of Ca2^+^-dependency LIME1 (*52*) further supported qualitative differences in signaling cascades. Both newly identified TCRs, KIF-sc1 and -sc2, differed from those opposite transcriptional patterns.

To distinguish TCR-intrinsic features from those potentially imprinted by previous antigen encounter or other patient-specific properties, we employed T cell populations from healthy donors transduced with defined neoTCRs and investigated functional patterns of these TCRs. This makes our approach especially interesting for the identification of transcriptomic reactivity patterns predicting functional TCR-capacity for ACT. Primarily, we focused on activation and inhibitory regulation centrally distinguishing SYTL4^S363F^-reactive from the other TCRs in our scRNAseq dataset. These analyses in neoTCR-tg T cells largely reflected the overall spectrum of our transcriptomic signatures strengthening TCR-inherence and transferability of the described patterns. Moreover, further distinction between KIF-sc2 (closer to KIF-P1 and -P2) and KIF-sc1 (closer to SYTL4^S363F^-reactive TCRs) illustrated unexpected heterogeneity in TCR-mediated reactivity for different clonotypes against one single peptide-HLA-complex (KIF2C^P13L^). Strong expression of activation markers and in particular the inhibitory receptors LAG-3 and PD-1 upon specific stimulation in vitro corroborated the distinction between burst-like, highly responsive but simultaneously strong inhibitory patterns (SYT-T1 and KIF-sc1) and otherwise moderate expression of activation markers (KIF-sc2 and -P2) with limited opposing inhibition. Our observation is in line with previous reports accounting a certain threshold of stimulation for the initiation of opposing inhibitory programs as a protective rheostat mechanism during early T cell activation (*53, 54*). Moreover, transferability of such qualities by gene transfer indicates a dominant structural or mechanic component of TCR-peptide-MHC complex assembly inherent to a defined TCR rather than imprinted differentiation responsible for these response patterns. Currently, the upregulation of inhibitory receptors is mostly linked to dysfunction and exhaustion (*14, 16, 21*) which are mainly understood as a result of chronic antigen encounter (*55, 56*). Other reports, however, argue for the initiation of T cell dysfunction early during tumorigenesis centrally determined by the quality of T cell stimulation and potential overstimulation (*57*). High levels of simultaneous activation and inhibitory regulation as well as induction of activation induced cell death observed as part of the burst-like activation pattern in our approach suggested TCR-driven hyperresponsivity, which, however, did not manifest in dysfunction upon first tumor challenge.

To understand potential TCR-dysfunction in the context of chronic stimulation, we investigated the persistence and resilience of T cells transgenic for KIF-sc1 and KIF-P2 representative for the two opposing response patterns recognizing the identical antigen in a rechallenge model. To mimic repeated antigen encounter we adapted our in vivo model and functionally investigated restimulated TILs from tumor explants from a xenograft NSG-mouse model in vitro as well as in vivo. In this setting, we detected significant functional impairment of KIF-sc1 with higher initial activation level and stronger opposing negative regulation upon antigen-specific restimulation compared to KIF-P2 with lower initial activation level. Overall, more moderate neoTCRs (especially KIF-P1 and -P2), with comparably high frequencies in the patient, transcriptomic patterns without negative regulation, slightly lower responsivity and functional avidity in vitro revealed potent in vivo-tumor rejection especially upon repeated stimulation (as shown for KIF- P2). From these findings we hypothesize, that 1. TCR-intrinsic features qualitatively determining activation have an enduring impact on the functional state and that 2. burst-like, highly responsive T cell reactivity patterns inducing high levels of inhibitory regulation are associated with functional impairment upon repeated stimulation.

These hypotheses certainly require further comparison with additional TCRs, also from other patients. The TCRs identified in Mel15 covered an only small range of comparably overall lower functional avidity compared to another recent publication which revealed high functional avidity as surrogate marker for “high-functionality” TCRs in their setting (*27*). Notably, even the slight differences between the TCRs compared in our study supported the notion of stronger upregulation of activatory and inhibitory signals on T cells expressing TCRs with higher functional avidity, as described by others (*27, 58*). Nevertheless, the substantial differences in maintained anti-tumor reactivity upon rechallenge, despite only such minor differences in functional avidity, suggest additional complexity of individual neoTCR activation patterns potentially associated to structural compounds, binding properties or inherent signaling differences. Moreover, T cell persistence and resilience upon repeated antigen challenge depicts a favorable TCR-quality requiring assessment for T cell engineering in ACT. Substantiated by other reports, moderate and potentially more stable rather than overly strong stimulation of T cells was associated with beneficial proliferation patterns and longevity of T cells (*44, 59, 60*). Meanwhile, another recent report underlined the importance of TCR-signal strength for development of T cell dysfunction by suggesting an intermediate level to maintain most potent anti-tumor efficacy over time (*26*).

In a synopsis, experimental outcomes like these and ours, have implications for T cell engineering and vaccination strategies (*61, 62*) currently focusing mainly on enhancing co-stimulatory receptor interactions (*63, 64*), reducing inhibitory signals (*65*) or TCR affinity maturation (*66, 67*). We show that individual TCR-intrinsic factors play a major role in determining T cell activation and sustained functionality in addition to p-HLA-complex density, antigen expression, co-signaling interactions and immunosuppressive factors in the TME. The question arising for future investigations therefore is, whether TCRs with qualitatively distinct activation profiles are necessary to complement each other in ACT. Whereas strongly activating TCRs might play a role in initial tumor debulking (under adequate ICI modulation), we hypothesize a substantial role for TCRs exhibiting more moderate stimulation patterns in sustained and resilient long-term tumor control.

## Material and Methods

### Primary patient material and cell lines

In line with the regulations and approval of the institutional review board (Ethics Commission, Faculty of Medicine, TU München) and in accordance with principles put forth in the Declaration of Helsinki, informed consent from all participants in this study was granted in written form.

The clinical course of melanoma patient Mel15 was previously described in detail (*28*). The identification of neoantigens resulting from somatic mutations (SYTL4^S363F^ and KIF2C^P13L^) by MS and in silico-prediction were previously reported (*12, 28, 29*). The PBMC sample of Mel15 used for single-cell-sequencing was selected based on previously confirmed reactivities within tested primary material at the specified time point (*28*), i.e. 966 days after first Ipilimumab application and 41 days after start of therapy with Pembrolizumab in a stage IV without evidence of disease.

PBMCs were isolated using density-gradient centrifugation (Ficoll-Paque) from either EDTA-anticoagulated blood of patient Mel15 (*28*), EDTA-anticoagulated blood or leukapheresis products from healthy donors. PBMCs were either immediately included in further downstream assays or stored in freezing medium (90% FCS and 10% DMSO) in liquid nitrogen. Feeder cells used in this study included pools of irradiated healthy-donor PBMCs.

Cell lines used in this study included as target cell lines: Mel15 lymphoblastoid cell line (LCLs) generated from Mel15 B cells by infection with Epstein-Barr Virus (EBV)-containing supernatant, U698M B cell lymphoma cell line (DSMZ cat# ACC-4) endogenously HLA-A03:01^+^ and HLA- B27:05^+^ as well as stably transduced with the mutated (mut mg) or wildtype (wt mg) tandem- minigene (*28*) and a fluorescent marker (Discosoma red fluorescent protein (dsRed) or green fluorescent protein (GFP)) and JJN3-B27 multiple myeloma cell line (DSMZ cat# ACC-541), endogenously HLA-A03^+^ and stably retrovirally transduced with HLA-B27 as described earlier (*12*). For retroviral transduction the embryonal kidney cell line 293Vec-RD114 (BioVec Pharma, Québec, Canada) stably expressing gag/pol and env was employed. For mouse experiments NS0-IL15 cells, kindly provided by S. R. Riddell in 2011, were used.

T cells were cultivated as previously reported (*12*). Target cell lines were cultivated in RPMI 1640 supplemented with 10% FCS, glutamine, non-essential amino acids, sodium pyruvate, and Penicillin/Streptomycin (Mel15 LCL, U698M, T2) or 40% DMEM + 40% IMDM supplemented with 20% FCS and Penicillin/Streptomycin (JJN3 B27). RD114 cells were cultivated in DMEM supplemented with 10% FCS and Penicillin/Streptomycin. Growth and morphology of cultivated cells were checked routinely. Absence of mycoplasma infection in cell lines and media was regularly confirmed by PCR or a cellular-based detection assay (PlasmoTest Mycoplasma Detection Kit).

### CD137 enrichment, rapid expansion and restimulation

To enrich PBMCs from patient Mel15 for KIF2C^P13L^- and SYTL4^S363F^-specific TCRs, we adapted our previously described method for identification of neoantigen-specific TCRs (*12, 28*). Both neoepitopes arose from a non-synonymous point mutation, resulting in naturally presented ligands on HLA-A03:01 for peptide KIF2C^P13L^ (amino acid sequence RLFLGLAIK) and HLA- B27:05 for SYTL4^S363F^ (GRIAFFLKY). Briefly summarized, PBMCs from Mel15 were cultivated in AIM-V supplemented with cytokines. After 24h, both neoepitope peptide ligands, KIF2C^P13L^ and SYTL4^S363F^ (0.1 μM) were added to the culture. Another 24h later, reactive T cells were separated using magnetic labelling and positive selection with the CD137 MicroBead Kit (Miltenyi). CD137^+^ enriched cells were then co-incubated with irradiated feeder cells in T cell medium (TCM) with supplements and expanded for eleven days.

After expansion, T cells were stimulated again with mutated KIF2C^P13L^ and SYTL4^S363F^ peptides using autologous antigen-presenting cells. Therefore, Mel15 LCLs were pulsed either with 0.1 µM KIF2C^P13L^ or SYTL4^S363F^ and irradiated with 30 Gy. Expanded T cells and irradiated LCLs were co-cultured at a ratio of 10:1 (T cells:LCLs) for 24 hours.

Interferon-gamma (IFN-γ) release of T cells was also assessed before and after enrichment using ELISpot assay as described before (*12*). Briefly, ELISpot plates were coated with IFN-γ capture antibody 1-DK1 (Mabtech) and incubated with cells. After removal of cells, anti-IFN-γ 7-B6-1 (biotinylated, Mabtech) as well as streptavidin-horseradish complex was added for visualization.

### CD8 isolation, scRNA-seq and scTCR-seq

CD8^+^ T cells were negatively isolated from the enriched, restimulated as well as an unstimulated Mel15-PBMC sample from the same timepoint using the Dynabeads™ Untouched™ Human CD8 T Cells Kit (Invitrogen). Single, alive (Propidium Iodode (PI)-negative) cells were sorted, 25 x 10^3^ cells from each sample were loaded onto one lane of a Chromium Next GEM Chip G (10x Genomics) and used for library prep using the Chromium next GEM Single Cell VDJ V1.1, Rev D) workflow (10x Genomics) as per company protocols. A high sensitivity dsDNA was used for quality control and analyzed on a Bioanalyzer 2100. Quantity of dsDNA was measured using a Qubit dsDNA HS kit (Life Technologies). Libraries were sequenced on an Illumina NovaSeq 6000 using read lengths of 26 + 8 + 0 + 91 for combined assessment of single cell RNA sequencing (scRNA-seq) and TCR sequencing (TCR-seq) information.

### Single-cell sequencing data bioinformatic analyses

Samples were converted from BCL to FASTQ using bcl2fastq (demultiplexed).

Raw paired-end sequencing files of the GEX and VDJ libraries were aligned to the human reference genome (refdata-gex-GRCh38-2020-A) and VDJ reference (refdata-cellranger-vdj- GRCh38-alts-ensembl-4.0.0) respectively, using 10x Genomics Cell Ranger (v4.0.0). Subsequently, we used the R package Seurat (v. 4.1.0) (*68*) to further analysis of the transcriptome- and TCR-based data. Only the genes detected in at least three cells were included in the raw counts matrix of the object. We retrieved only the cells containing at least 200 genes and fewer than 6000 genes. To avoid possible dead cells contamination, we excluded the cells with a fraction of mitochondrial genes higher than 18%. In the next step, the corresponding TCR data was added to the meta.data slot of the Seurat object. Raw gene counts were log-normalized, and variable features were detected with the vst method. Subsequently, canonical correlation analysis (CCA) integration was used to leverage the batch effects between two experimental setups combined in one Seurat object. After that, we newly determined the variable features using the integrated assay and scaled the expression matrix with regression on the number of UMIs and fraction of mitochondrial genes per cell. Unbiased calculation of k-nearest neighbors was done, and using UMAP, neighborhood graph and embedding were generated. After the UMAP construction, we retrieved only the cells containing the TCR information and clonotypes expressing more than one alpha or beta chain were removed. Previously identified neoTCRs from our index patient (*28*) were detected using their CDR3 region, and corresponding clonotypes in our samples were assigned to the respective TCR group. The final cell numbers in our linked TCR-transcriptome data set were 5764 cells in the unstimulated and 6007 cells in the restimulated sample. The FindAllMarkers function was used to calculate differentially expressed genes in each cluster. The corresponding upregulated genes were retrieved for the subsequent pathway enrichment analysis using the enrichR (v. 3.0) R package. Seurat clusters were annotated manually by analyzing the expression of upregulated genes on the UMAP. The gathered signature expression score was generated by using AddModuleScore function. Subsequently, the Seurat object was converted into .h5ad format, and the pseudotime score with corresponding diffusion maps was generated using the scanpy library implemented in Python (*69*). For pseudotime score calculation, cluster 1_CCR7 (most naїve) was set as a starting point.

For the differential gene expression analysis between the TCR groups, we used the FindMarkers function of the Seurat package by plotting the results using the ggplot2 (v. 3.3.5) R package.

### V(D)J analysis and selection of TCRs for TCR transduction

For subsequent TCR selection a meta data .csv was exported after initial QC (s. above). Only clonotypes expressing exactly one productive alpha and one beta chain were considered to allow for precise identification of TCRs. The total number of this refined TCR set was 4182 in the unstimulated and 4913 in the restimulated sample. To select new neoTCRs, clonotypes that had previously been identified were excluded and the frequencies of remaining clonotypes were compared. We considered two metrics: highest fold change of TCR frequency before and after stimulation as well as greatest absolute frequency of clonotypes in the restimulated sample. We selected four new TCRs for investigation of specificity and functionality, two of them demonstrated specificity for KIF2C^P13L^, later termed TCR KIF-sc1 and -sc2.

### Engineering of KIF-sc1 and -sc2 TCRs

α- and β-chain-sequences of clonotypes identified as potential neoantigen-reactive TCRs were submitted to IMGT to obtain comprehensive information on respective V-D-J sequences (https://www.imgt.org/IMGT_vquest/vquest). Full-length TCR sequences were reconstructed using Ensembl database and subsequently in silico optimized throughout insertion of a cysteine bridge, murinization of the constant region and codon optimization (*70–72*). β- and α-chain were linked by a P2A element and tandem gene products were synthesized (BioCat). Each TCR candidate was cloned into MP71 retroviral vector and subsequently used for transduction into healthy donor T cells.

### Retroviral transduction of healthy donor CD8^+^ T cells with neoTCRs

CD8^+^ T cells used for transduction were obtained by magnetic negative selection from healthy donor-derived PBMCs (EasySep™ Human CD8^+^ T Cell Isolation Kit, Stemcell) and activated for 48h with 30 U/ml human IL-2 and anti-CD3-anti-CD28-beads (Dynabeads™ human T-Activator CD3/CD28, ThermoFisher). Retroviral packaging cells RD114 were seeded to reach a confluency of 60% on the day of transfection and subsequently transfected with plasmids containing the neoTCR-α- and -β-chain-sequences using TransIT®-293 (MirusBio). Transfected cells were incubated for 48 h and supernatants subsequently filtered and used for spin infection of activated CD8^+^ T cells. Transduced T cells were cultivated with IL-7 and IL-15 for 10 days as described before (*12*). Transduction efficacies were determined via fluorescence activated cell sorting (FACS) staining with TCRmu antibody (anti-mouse TCR-β-chain, FITC, BD Biosciences) against the murine-β-chain of engineered TCR-constructs in comparison to non-transduced T cell populations.

### In vitro assessment of reactivity and activation patterns in TCR-tg T cells

The subsequently described functional and phenotypic aspects were assessed within co-culture settings using transgenic CD8^+^ T cells from different healthy donors and different target cells. Cell lines were either transgenic for the tandem minigene (mut mg versus wt mg) or pulsed with different concentrations of peptides KIF2C^P13L^ and SYTL4^S363F^, their wildtype form or peptide derivates containing single amino acid substitutions with alanine and threonine at all possible positions as described before (*28*). FACS- as well as ELISA-based readout was performed at different timepoints after co-culture setup as indicated. In selected experiments, varying transduction efficiencies (between donors and transductions) were equalized by diluting to the lowest rate per assay with a minimum at 10% of TCRmu^+^ cells with non-transduced CD8^+^ T cells obtained from the same donor. TCR-tg TCRmu^+^ T cells were considered effector cells for all E:T-ratios.

### Extra- and intracellular FACS staining

FACS staining was performed in FACS buffer (PBS with 1% FCS and 2mM EDTA) in 96well-u-bottom plates. Cells from in vitro co-cultures or tumor lysates were washed in FACS buffer and for in vitro co-cultures experimental triplicates were pooled prior to staining. Unspecific binding sites were blocked with 30% human serum in FACS buffer for 20 min at 4°C before extracellular (EC) staining with diverse antibodies diluted in FACS buffer at 4°C for 30 min. Live/dead stains were either directly added to the EC-antibody mix (Hoechst, thermofisher) or added directly prior to measurement (PI, 7-AAD).

In addition to EC-staining subsequent intracellular (IC)-staining was performed for several analyses. Prior to EC-staining, fixable live-dead stain (Zombie UV, biolegend) was stained in PBS. After EC-staining, cells were washed and fixed (Fixation buffer, biolegend) for 20 min at RT (protected from light). Afterwards, perm buffer (biolegend) diluted in deionized water was used for permeabilization according to manufacturer’s protocol. IC-staining antibody-mix in perm buffer was added afterwards for 40 min at RT (protected from light), followed by further washing steps.

FACS antibodies used for analyses: aCD137-PE, aCD137-APC and aCD137-APC-Cy7 (all biolegend, clone 4B4-1), anti-murine TCR-β-FITC (BD Biosciences, H57-597), aPD-1-BV785 (biolegend, EH12.2H7), aLAG-3-BV650 (biolegend, 11C3C65), aCD3-AF700 (biolegend, UCHT1), aCD8-PerCP (biolegend, SK1), aIFN-γ-APC (biolegend, 4S.B3), aGzmB-PE (biolegend, QA16A02).

Sample analysis was performed at an LSRII or LSR Fortessa (BD Biosciences). FACS data was analyzed using Flow Jo_v10.8.1.

### Activation induced cell death (AICD) assessment: Annexin-V staining

Cells stained extracellularly with FACS antibodies were stained in Annexin-V binding buffer diluted in water (thermofisher) with AnnexinV (APC, biolegend) and PI for 20 min at RT prior to analysis.

### Proliferation assessment: Cell Trace Violet (CTV)-staining

TCR-tg CD8^+^ T cells from three healthy donors were labelled with CTV Dye (thermofisher) according to manufacturer’s guidelines. On day 4 of co-culture, cells were stained extracellularly with FACS antibodies and afterwards the percentage of TCRmu^+^ T cells per division was determined via flow cytometric readout.

### In vivo tumor rejection potential in a xenograft model

NOD.CG-Prkdcscid IL2rgtm1Wjl/SzJ (NSG; The Jackson Laboratory) were maintained according to the institutional guidelines and approval of local authorities. A xenograft murine model was established as previously described (*73, 74*). Animal well-being was assessed daily and tumor growth was monitored in vivo by external measurements with digital caliper.

### Tumor rejection potential of TCR-tg T cells

The capacity of primary tumor control was assessed as described before (*28*). Briefly, NSG mice at the age of six to nineteen weeks were subcutaneously injected with U698M-mut mg cells (10x10^6^ cells/flank). As tumors reached an area of ca. 20 mm^2^, T cells transduced with TCRs KIF- P2, KIF-sc1, KIF-sc2, SYT-T1 or T cells transduced with an irrelevant TCR (2.5D6 targeting MPO (*73*)) were injected intravenously. 2x10^7^ transduced T cells (3.2x10^7^ absolute T cells including non-transduced cells) were administered to 6 mice per group (n = 6) in two injections on two subsequent days. Male and female animals as well as animals of different age were distributed evenly across all treatment groups.

### Rechallenge model: Tumor rejection potential of TIL products generated from transgenic T cells

For the generation of TIL products, female NSG mice at the age of seven weeks were subcutaneously injected with U698M-mut mg cells (10x10^6^ cells). As tumors reached an area of 20 mm^2^, T cells transduced with TCRs KIF-P2 and KIF-sc1 were injected intravenously. 8x10^6^ transduced T cells (in total 11x10^6^ including non-transduced cells) were administered to five mice per group (n = 5) at equalized transduction rates of 70 % for both groups. Tumor growth kinetics were monitored daily for 5 days with digital caliper. On day 5, before tumor regression was measurable, animals were sacrificed, tumors as well as spleen explanted and passed through a cell strainer (100 μm). Partly, tumor material was used for immediate downstream applications. Further parts of the tumor material were cultivated with irradiated (70 Gy) feeder cells, 1000 U/ml human IL-2 and 30 ng/ml anti-CD3 antibody (OKT3) in TCM for 21 days. IL-2 was supplemented on days 7, 11 and 15 (300 U/ml). Efficacy of TIL generation was assessed by FACS staining (CD8 and TCRmu) and TILs of those two mice with the highest rate of TCRmu^+^ CD8-TILs per TCR pooled to reach equal transduction rates for all subsequent experiments (55%).

We injected 5x10^6^ transgenic T cells (transduction rate of 55%) into five U698M-tumor bearing mice (equal distribution of male and female, age between 7 and 15 weeks) per group. We applied KIF-P2 and KIF-sc1 TCR-tg T cells each for the TIL-P-conditions (injected on day 21 after tumor explant; 43 days after blood donation) as well as a new batch of transgenic T cells from the same donor (17 days after blood donation). We compared this with 2.5D6-tg T cells as negative control. Tumor growth kinetics were monitored daily by blinded measurement regarding T cell condition for 30 days with digital caliper until experiment endpoint criteria were reached.

### Statistics

Significance of differences between TCR products on the day of explant or of TIL reinfusion were investigated by one-way analysis of variance (ANOVA) and Tukey’s multiple comparison test. Regarding the rechallenge model, differences in tumor growth were calculated for the tumor area on day 17 with ordinary one-way ANOVA and Tukey’s multiple comparison test. Statistical comparison of survival was performed using the Mantel-Cox test. Statistical analyses were performed with GraphPad Prism V.9.3.1 software.

## Supporting information

Fuechsl & Untch et al_Supplementary Figures and Table legends

Fuechsl & Untch et al_Supplementary Tables

## Acknowledgements

The authors thank the patient for participating in the study and his continuous support. We also thank Stefanie Stein for excellent technical support. We thank Michael Hiltensperger for critical reading of the manuscript.

## Funding

EIT Health 19638 (AMK)

Deutsche Forschungsgemeinschaft (DFG, German Research Foundation) –SFB824 (C10 AMK) and SFB- TRR 338/1 2021 –452881907 (A03 AMK/ A01 DHB)

Else Kröner-Fresenius-Stiftung, doctoral program „Translationale Medizin“ (JU)

## Author Contributions

Conceptualization: FF, JU, EBr, AMK

Methodology: FF, JU, EBr, VK, SJ, CV, NdAK, DG, EdB and AMK Experimental Work: FF, JU, EBr, SJ, CV, DG

Data analysis: FF, JU, EBr, VK, NdAK, EdB and AMK Visualization: FF, JU, EBr, VK, AMK

Funding acquisition: DHB, AMK Project administration: EBr, AMK Supervision: RR, DHB, EBe, EBr, AMK

Writing – original draft: FF, JU, EBr, AMK Writing – review & editing: all authors

## Ethics approval

Informed consent of all healthy donors and the patient was obtained following requirements of the institutional review board (Ethics Commission, Faculty of Medicine, TUM) and in accordance with the principles of the Declaration of Helsinki.

