## Supplementary material for "High-resolution profiling of neoantigen-specific T cell receptor activation signatures links moderate stimulation patterns to resilience and sustained tumor control": Fuechsl & Untch et al_Supplementary Figures and Table legends

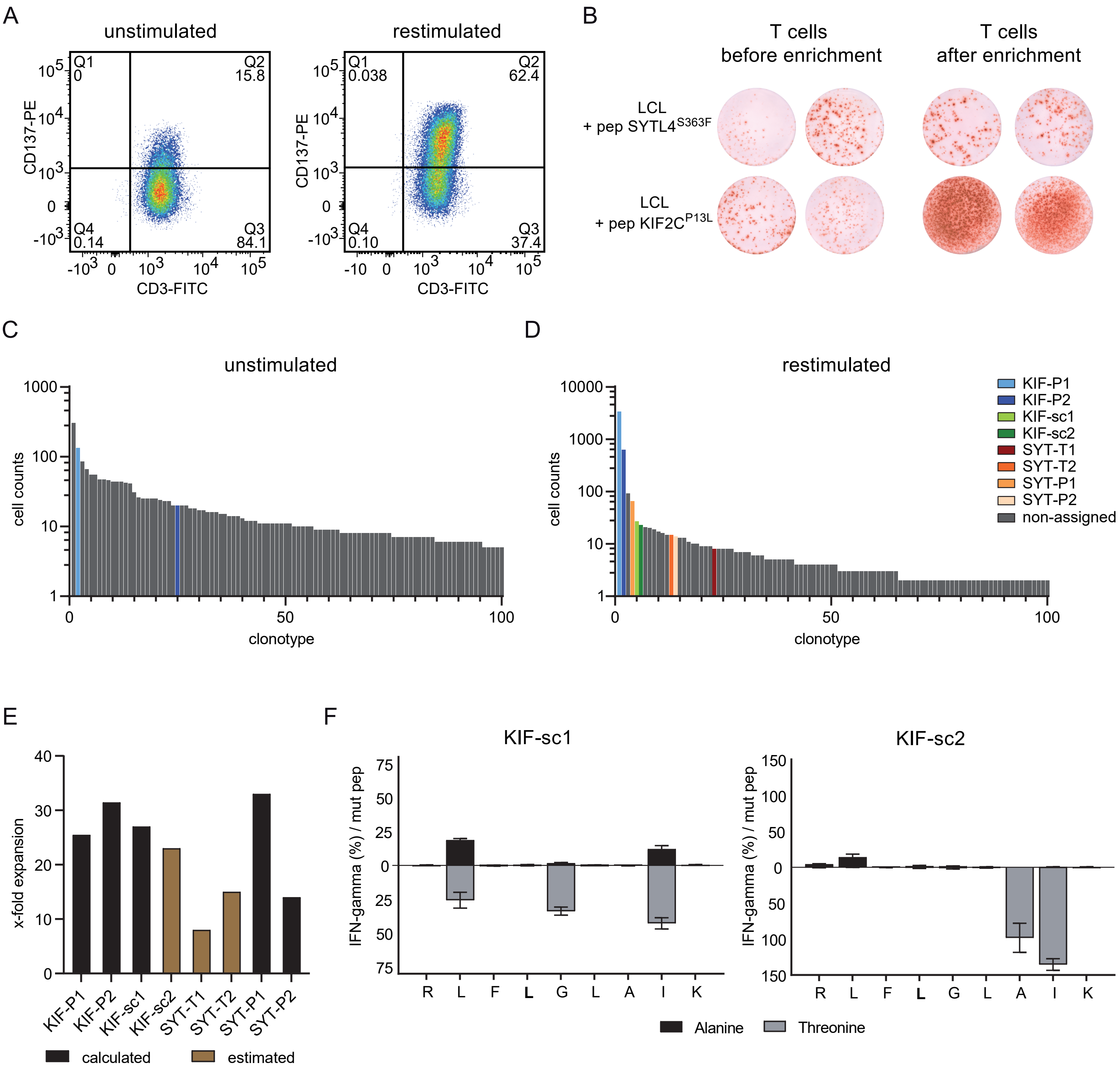


**Supplementary Figure 1**

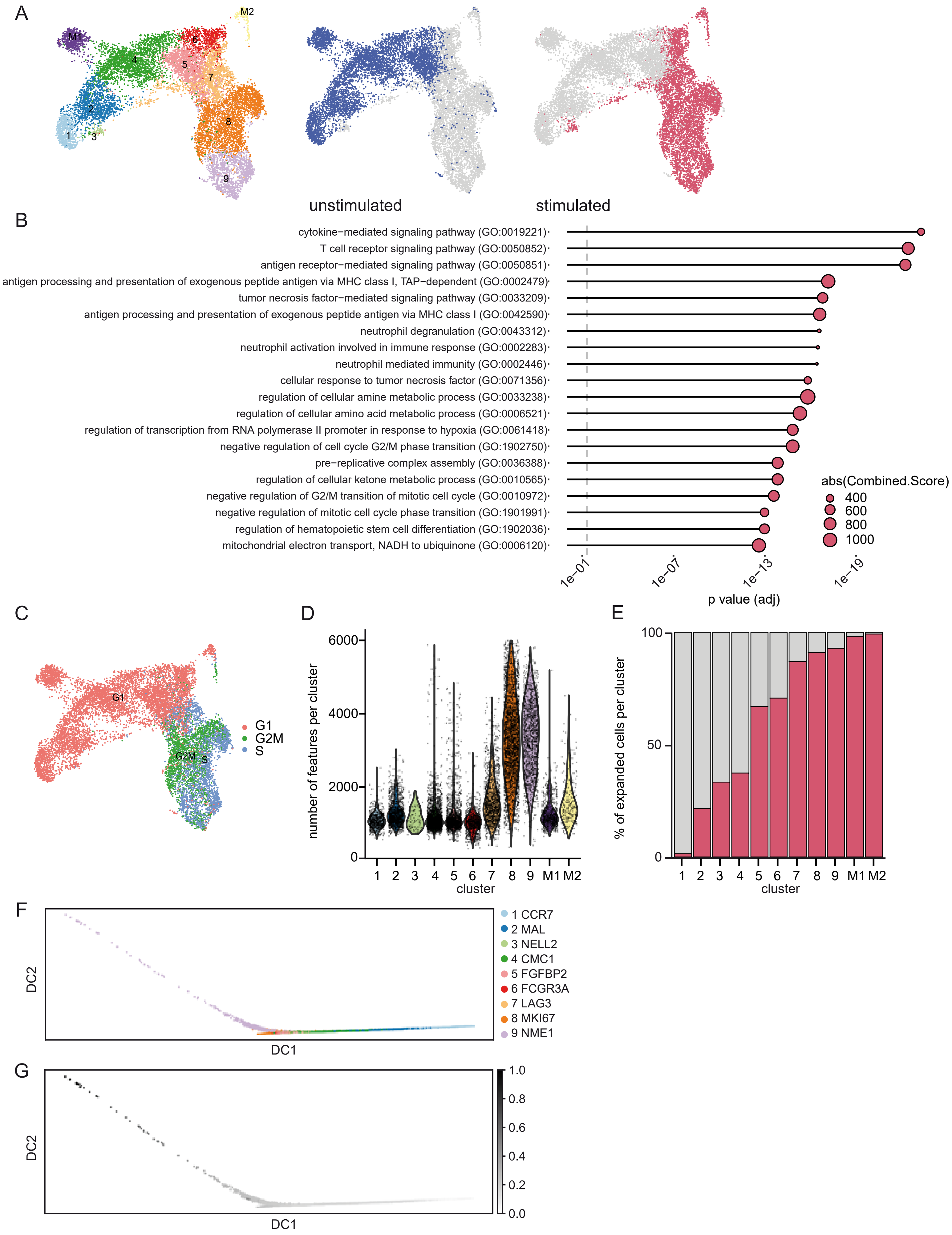


**Supplementary Figure 2**

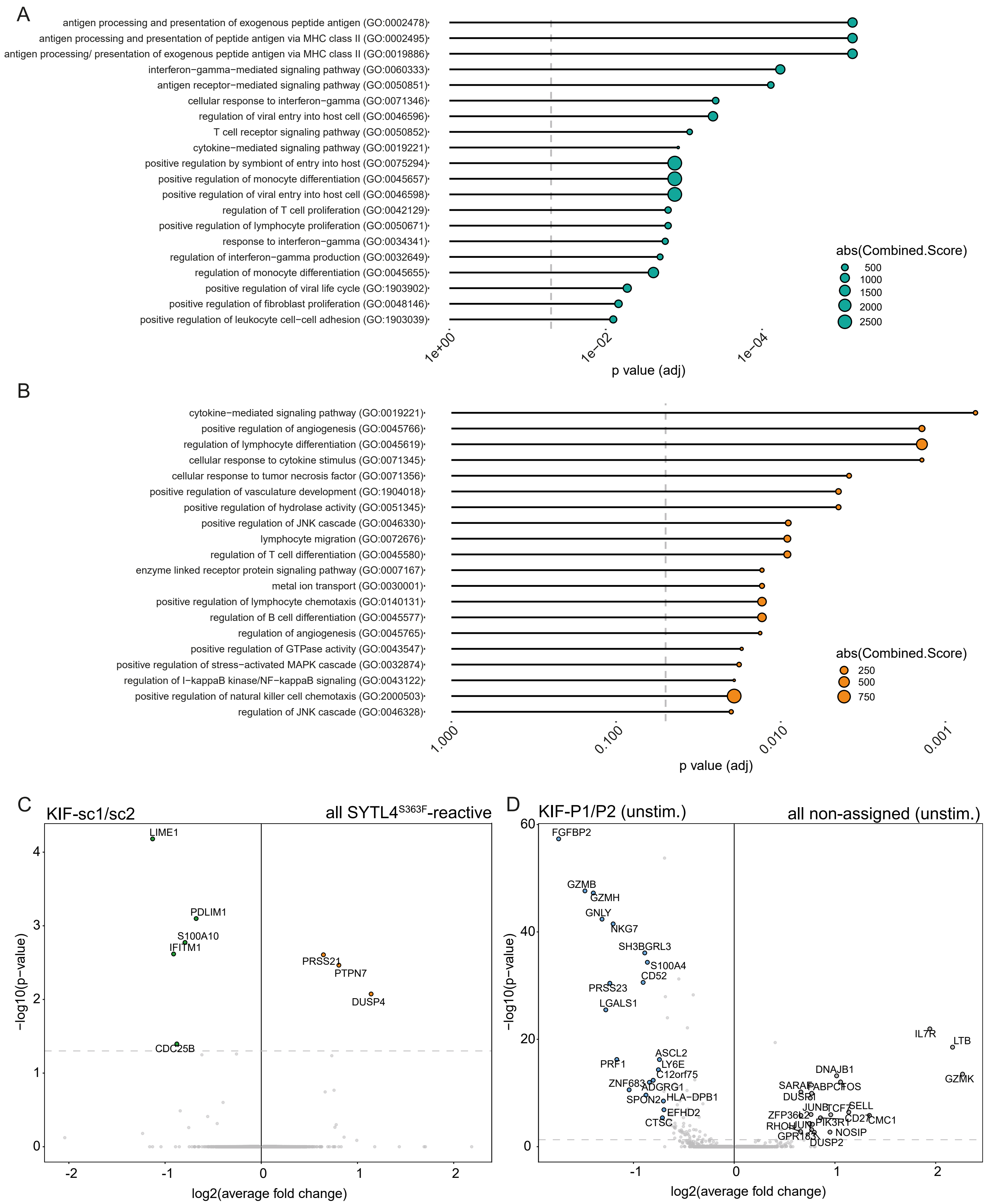


**Supplementary Figure 3**

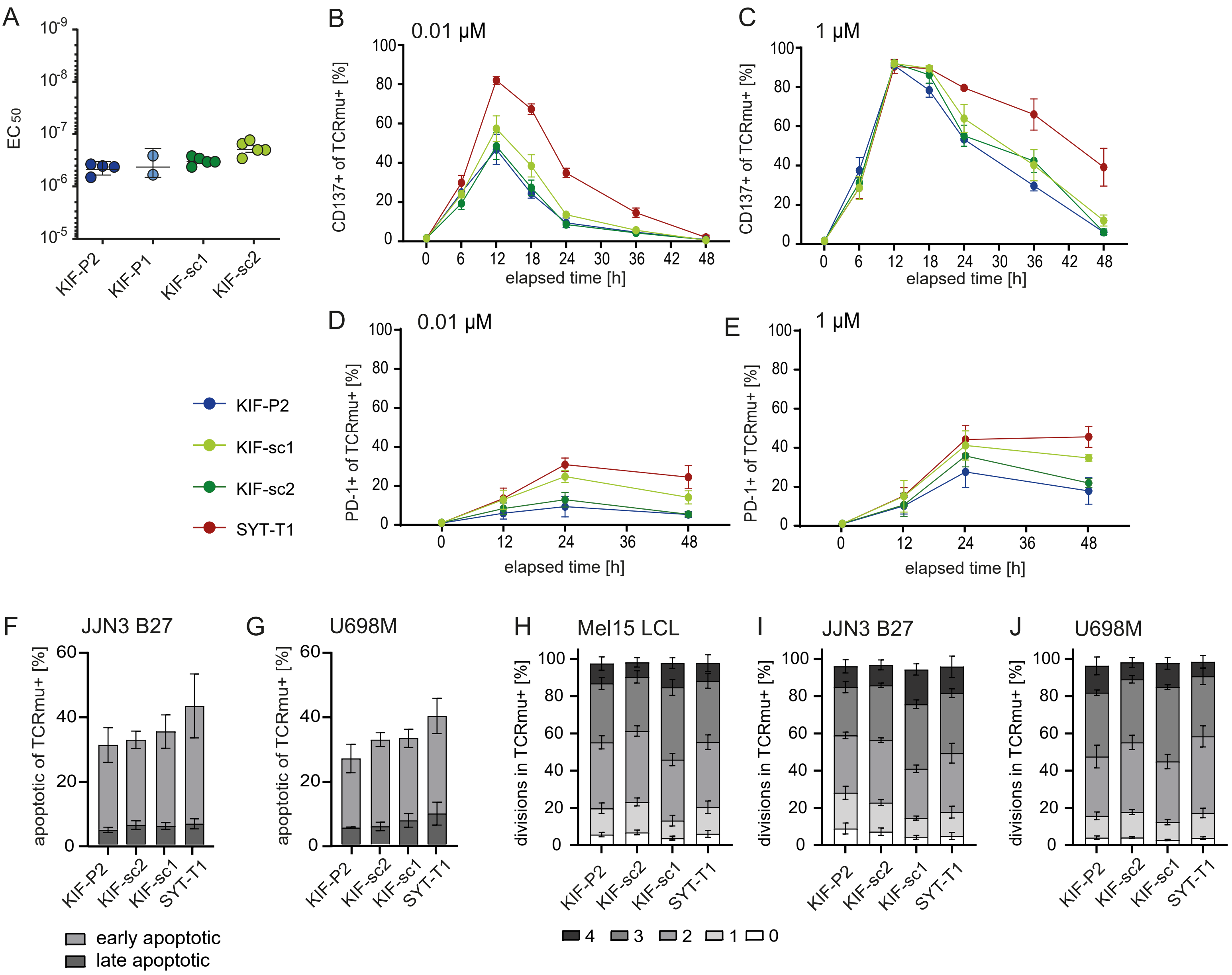


**Supplementary Figure 4**

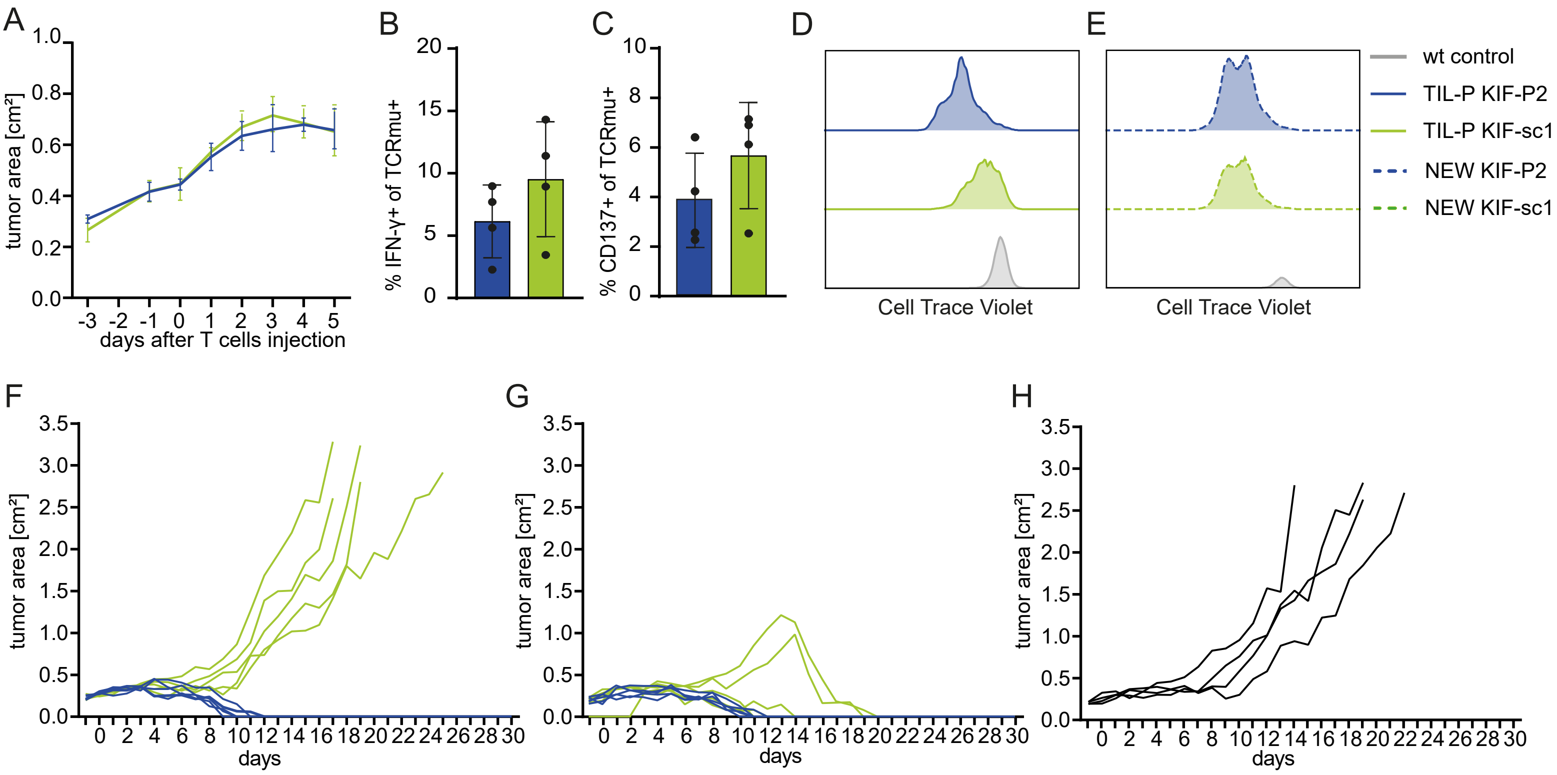


**Supplementary Figure 5**

### Supplemental Information

**Supplementary Figure 1. CD137-based enrichment of neoepitope-stimulated PBMCs shapes the TCR repertoire and leads to the identification of additional KIF2C$^{P13L}$-specific TCR.** A, Flow-cytometric assessment of CD137 surface expression measured before and after CD137-specific enrichment of SYTL4$^{S363F}$- and KIF2C$^{P13L}$-peptide stimulated PBMCs from patient Mel15. B, Comparison of spot forming units (SFU) in IFN-γ ELISpot of Mel15 samples before and after CD137-enrichment. T cells were co-cultured with LCL pulsed with the indicated peptide. C, D, Frequencies of the top 100 prevalent TCR clonotypes within the unstimulated (C) and stimulated and enriched (D) scRNA-seq dataset. E, Bar plot showing the fold expansion of neoTCR clonotypes comparing frequency before and after stimulation. For neoTCRs not detected in the unstimulated sample, an estimated ratio was calculated using a theoretical frequency of x = 1 read in the unstimulated sample. F, Amino acid recognition motif of neoTCRs KIF-sc1 and -sc2 determined by alanine/threonine scan. TCR-tg T cells were co-incubated with target cells pulsed with peptides containing an amino acid substitution with alanine (upper graph) or threonine (lower graph) at the indicated position. Percentage of recognition was calculated as percentage of IFN-γ response in relation to IFN-γ response against target cells pulsed with the original mutated peptide KIF2C$^{P13L}$.

**Supplementary Figure 2. Expression profiles of known and unknown TCR clonotypes were compared as assessed by scTCR-/scRNA-sequencing.** A, UMAPs of 5764 unstimulated (blue, middle graph) and 6007 restimulated (red, right graph), enriched (single alive) CD8$^{+}$ T cells next to the UMAP with all identified clusters (left graph). B, Top 20 gene ontology biological process (GO:BP) terms of the pathway enrichment analysis of the 7_LAG3 differential gene expression signature. C, UMAP indicating transcriptomic analysis of cell cycle phase of each cell. Color code indicating G1 (red), G2M (green) and S (blue) phase. D, Violin plot showing number of identified genes per cell within the clusters. E, Bar plot indicating percentage of expanded cells within each cluster (clonotypes with two or more cells count as expanded). F, Diffusion map using the cells of the 9 clusters excluding MAIT cells. G, Pseudotime scores plotted on the diffusion map. Starting point was set on the most naïve cluster (1_CCR7).

**Supplementary Figure 3. Comparison of transcriptomic activation signatures of neoTCR-groups in the repertoire of patient Mel15.** A, Top 20 GO:BP terms of the pathway enrichment analysis of KIF2C$^{P13L}$-specific TCRs. B, Top 20 GO:BP terms of the pathway enrichment analysis of SYTL4$^{S363F}$-specific TCRs. C, Volcano plot comparing fold changes of transcriptomic expression in all SYTL4$^{S363F}$-specific TCRs against KIF-sc1/-sc2 TCRs. D, Volcano plot showing differential transcriptomic gene expression of unstimulated TCRs KIF-P1 and -P2 in comparison to all non-assigned clonotypes within the unstimulated T cell fraction.

**Supplementary Figure 4. Co-cultures with different cell lines and activation markers reflect distinct activation signatures for CD8 neoTCR-tg T cells in vitro.** A, Assessment of $EC_{50}$-values for all KIF2C-reactive neoTCRs of IFN-γ-secretion measured by ELISA from co cultures with peptide-pulsed T2 cells for several different donors without equalizing $TCRmu^{+}$-transduction rates. E:T = 10.000 T cells : 10.000 tumor cells. B-E, EC FACS staining of CD137 at different timepoints after co-culture setup with Mel15 LCL peptide-pulsed target cells (B, C; E:T = 1:1 (10.000 tg T cells : 10.000 tumor cells)) and of PD-1 after co-culture setup with JJN3 B27 peptide-pulsed target cells (D, E; E:T = 1:1 (15.000 tg T cells : 15.000 tumor cells)). A weak (0.01 µM for peptide pulsing; B, D) versus a strong (1 µM for peptide pulsing; C, E) stimulus are compared. F, G, Annexin-V/PI-staining was employed for detection of activation induced cell death (AICD) upon strong stimulation with 1µM mut-peptide pulsed JJN3-B27 (F) or mut mg-tg U698M target cells (G) after 20h of co culture (early apoptotic = $AnnexinV^{+}PI^{-}$, late apoptotic = $AnnexinV^{+}PI^{+}$). E:T = 1:1 (15.000 tg T cells : 15.000 tumor cells). H-J, CTV-labelling prior to co-culture was used to detect the frequency of cells per number of divisions (0 to 4 divisions) after 4 days of co-culture. Frequencies of divisions of all $TCRmu^{+}$ cells are shown for co cultures with Mel15 LCLs (H), JJN3-B27 (I) or mut mg-tg U698M target cells (J). For B-J technical triplicates of one donor were pooled prior to staining; the Mean and SD for biological replicates from three different human donors are shown if not stated otherwise.

**Supplementary Figure 5. Moderate activation pattern of KIF-P2-tg T cells is reflected in tumor-derived T cells and associated with more sustained functionality upon rechallenge.** A, Tumor growth kinetics are displayed as tumor area (in $cm^2$) for mut mg-U698M-tumor-bearing NSG-mice injected with $8x10^6$KIF-P2- (blue) versus -sc1-tg (green) T cells on day 0 of the TIL generation experiment depicted in Figure 5A (n=5). Mean values and SDs for each group of mice display tumor growth dynamics (n=5). Mice were sacrificed on day 5 after T cell injection for further analysis and TIL-generation. B, C, FACS staining of IFN-γ (IC; B) and CD137 (EC; C) on $CD8^{+}CD3^{+}/TCRmu^{+}$ T cells (KIF-P2 in blue, KIF-sc1 in green) harvested from mut mg-U698M-tumor explants upon sacrifice of mice on day 5 after T cell injection (n=4; only mice with sufficient numbers of detected $CD3^{+}CD8^{+}$ T cells in FACS analysis (>30) were included). The Mean and SD of four mice per group are depicted. D, E, CTV-labelled TIL-P (D) and NEW (E) TCR-tg T cells were analyzed on day 4 after co culture initiation with mut mg-U698M tumor cells in comparison to a representative control pulsed with wt mg-U698M tumor. E:T = 1:1 (50.000 tg T cells : 50.000 tumor cells). Triplicates were pooled prior to FACS analysis. F-H, Individual tumor growth curves per mouse of the experiment shown in Figure 5H are depicted for TIL-P-KIF-P2 (blue) and TIL-P-KIF-sc1 (green) (F), NEW-KIF-P2 (blue) and NEW-KIF-sc1 (green) (G) as well as 2.5D6 (black) (H).

**Supplementary Table 1.** Single cell characteristics extracted from sequencing data after QC.

**Supplementary Table 2.** NeoTCR clonotypes with specificity for KIF2C$^{P13L}$ and SYTL4$^{S363F}$ as well as TCR frequency before and after enrichment in the linked scTCR and transcriptome data set.

**Supplementary Table 3.** Recognition motif of neoTCRs extracted from alanine/threonine scan. Parts of this data have already been published (*28*) and were listed here next to the novel motifs of KIF-sc1 and -sc2 for direct comparison.

**Supplementary Table 4.** Percentage of each TCR contribution to the sum of all detected KIF2C$^{P13L}$- and SYTL4$^{S363F}$-specific T cells within $M_{Int}$, $M_{Int}$-LN1, $M_{Int}$-LN2, $M_{Lung}$ and $M_{Lung}$-LN. Parts of this dataset were published before, i.e. for the previously identified TCRs KIF-P1, -P2, SYT-T1, -T2, -P1, -P2 (*28*) and are listed here for embedding the newly identified TCRs in the entire setting.

**Supplementary Table 5.** $EC_{50}$ values of neoTCRs as a measure for functional avidity. Data for TCRs SYT-T1, -T2, -P1, -P2, KIF-P1 and -P2 were pooled from previous experiments (*28*) and listed next to data retrieved for KIF-sc1 and -sc2 as depicted in Figure S4A.
